# Predicting biological age based on the BBMRI-NL ^1^H-NMR metabolomics repository

**DOI:** 10.1101/632919

**Authors:** E.B. van den Akker, S. Trompet, J.J.H. Barkey Wolf, M. Beekman, H.E.D. Suchiman, J. Deelen, F.W. Asselbergs, BBMRI-NL, E. Boersma, D. Cats, P.M. Elders, J.M. Geleijnse, M.A. Ikram, M. Kloppenburg, H. Mei, I. Meulenbelt, S.P. Mooijaart, R.G.H.H. Nelissen, M.G. Netea, B.W.J.H. Penninx, M. Slofstra, C.D.A. Stehouwer, M.A. Swertz, C.E. Teunissen, G.M. Terwindt, L.M. ‘t Hart, A.M.J.M. van den Maagdenberg, P. van der Harst, I.C.C. van der Horst, C.J.H. van der Kallen, M.M.J. van Greevenbroek, W.E. van Spil, C. Wijmenga, A. Zhernakova, A.H. Zwinderman, N. Sattar, J.W. Jukema, C.M. van Duijn, D.I. Boomsma, M.J.T. Reinders, P. Eline Slagboom

## Abstract

The blood metabolome incorporates cues from the environment as well as the host’s genetic background, potentially offering a holistic view of an individual’s health status. We have compiled a vast resource of ^1^H-NMR metabolomics and phenotypic data encompassing over 25,000 samples derived from 26 community and hospital-based cohorts. Using this resource, we constructed a metabolomics-based age predictor (metaboAge) to calculate an individual’s biological age. Exploration in independent cohorts demonstrates that being judged older by one’s metabolome, as compared to one’s chronological age, confers an increased risk on future cardiovascular disease, mortality and functionality in older individuals. A web-based tool for calculating metaboAge (metaboage.researchlumc.nl) allows easy incorporation in other epidemiological studies. Access to data can be requested at bbmri.nl/samples-images-data. In summary, we present a vast resource of metabolomics data and illustrate its merit by constructing a metabolomics-based score for biological age that captures aspects of current and future cardio-metabolic health.

## MAIN

Chronological age is an important risk factor for virtually all types of common disease, including diabetes mellitus type 2, cardiovascular disease and many forms of cancer [1]. Moreover, chronological age is often employed as an important criterion on which clinical treatment decisions in older adults are based. Yet, especially in the elderly, chronological age is a poor representative of an individual’s intrinsic biological age, including the susceptibility to disease and resilience to treatment [2]. Hence, novel biomarkers are required that give additional information about the disparity between chronological and biological age, i.e. whether individuals are biologically older and potentially more vulnerable than their peers.

A range of multi-marker algorithms has been developed to serve as indicators of biological age. Examples are those based on physiological deterioration of organ systems from the second [3] or third [4] decade onwards, or those based on combined health deficits in later life, the so-called ‘frailty’ indices [5,6]. Others have exploited large quantities of highly standardized molecular data, e.g. DNA methylation data, to train so-called ‘clock’ algorithms [7-9] that allow one to calculate an omics-based age. The difference between an individual’s actual chronological age and the estimated ‘methylation age’ was for instance shown to associate with mortality [10]. Interestingly, when compared, each of these omics-based biological age indicators appeared to mark unique aspects of ageing [11,12], giving ample incentive for the development of other, possibly complementary omics-based indicators of biological age. However, the requisite of a large body of data initially required for training similar omics age predictors is currently limiting the application of other molecular domains.

## RESULTS

### The BBMRI-NL resource

We present a novel, well standardized ^1^H-NMR blood-based metabolomics data set encompassing over 25,000 samples collected by the Dutch Biobanking and BioMolecular resources and Research Infrastructure derived from 26 community and hospital based cohorts (**Figure 1 and Table S2,** available upon request at BBMRI-NL: bbmri.nl/samples-images-data, along with routine clinical variables). We have employed these data to construct a metabolomics-based clock (predictions made available as web resource (metaboage.reasearchlumc.nl, see **Online Methods** for instructions), and show that the difference between chronological age and ‘metabolomic age’ captures aspects of cardio-metabolic health.

**FIGURE 1:**
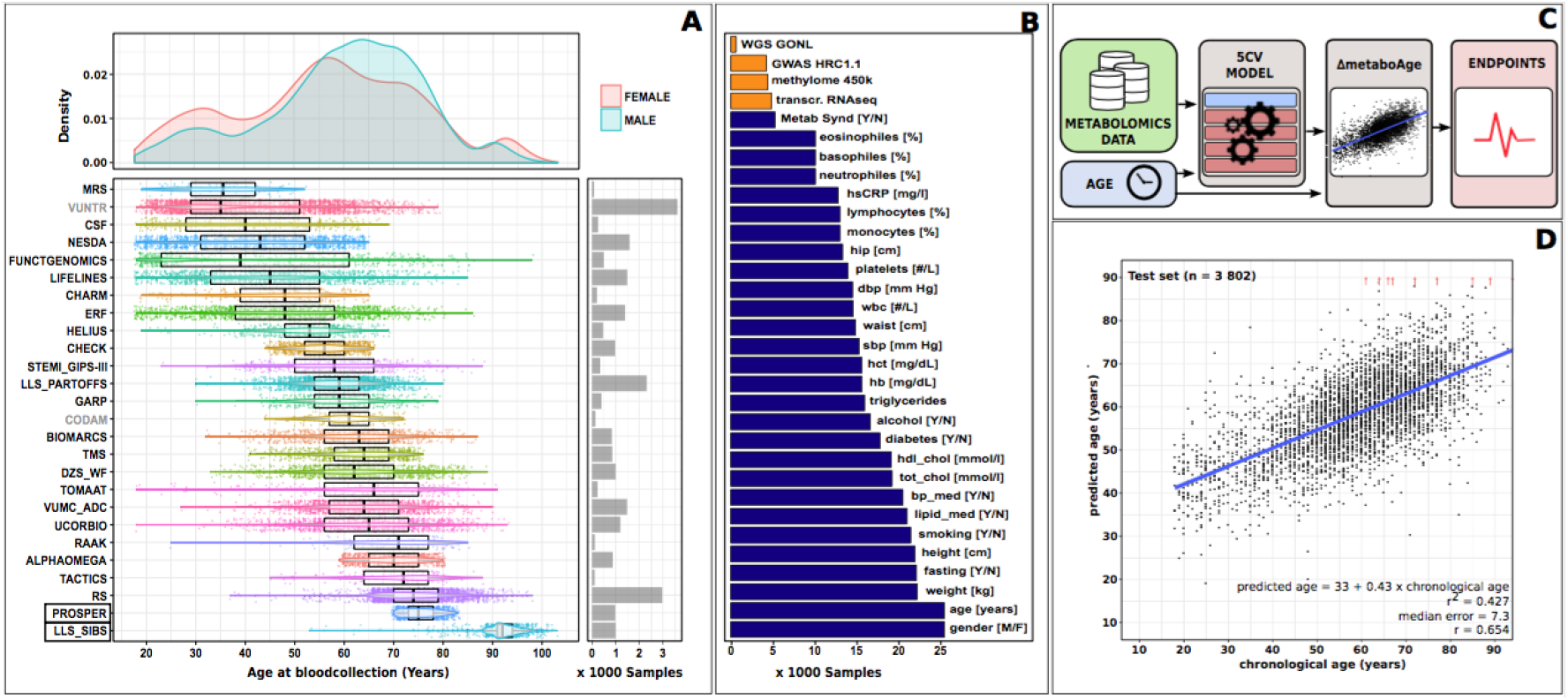
BBMRI-NL a vast ^1^H-NMR metabolomics resource enabling approaches for personalized medicine. **A:** Cohorts in the Dutch Biobanking and Biomolecular resources Research Infrastructure (BBMRI-NL), totaling 25,253 samples display interlinked age distributions robustly covering the complete adult lifespan from 18 till 85 years. While, VUNTR and CODAM (grey) were omitted for training the age predictor due to incomplete data (**Online Methods**), LLS_SIBS and PROSPER (boxed) were held out to independently evaluate the merit of age predictions as surrogate biomarkers for clinical endpoints. **B:** Additional omics data (orange) and phenotypic variables (blue) available within the BBMRI-NL resource. **C:** Flow chart of the analyses: a predictor for chronological age is trained on BBMRI-NL metabolomics data. The age-independent part of differences between predicted age and chronological age, termed ΔmetaboAge, is associated with endpoints. **D:** 5-Fold-Cross Validation (5FCV) is performed to assess the accuracy of the age predictor. Predictions on the test set of a representative fold are depicted, with ΔmetaboAge exemplified in orange.

### Deriving a metabolomics-based score for biological age

A metabolomics predictor for chronological age was trained and evaluated (see **DocS1**) employing 56 out of 226 most reliable and independent [13] metabolomic variables (see **Table S3 and DocS2**), derived from 24 cohorts (**Figure 1**). Two biobanks missing a metabolomic variable were omitted (see **Online Material**). In addition, PROSPER and LLS_SIBS were left out from training the metabolomic age predictor and used to independently explore the predictive value of the obtained indicator of biological age. With use of the data of the remaining 22 biobanks comprising 18,716 samples (9,680 males and 10,036 females), a linear model was trained with the 56 metabolomic variables to estimate chronological age (see **Online Material**). A 5-Fold-Cross-Validation (5FCV, see **Online Material**) scheme was employed for randomly splitting the data in training (80%, 15,208 samples) and test (20%, 3,802 samples) sets for an unbiased training and evaluation of the models. The age-independent part of the difference between the estimated metabolomic age and chronological age (**Figure 1C**), hereafter referred to as ΔmetaboAge, may reflect for each individual the disparity between their biological and chronological age. Consequently, a high ΔmetaboAge indicates a relatively ‘old’ blood metabolome for a given chronological age.

### Associations of metaboAge with cardio-metabolic risk factors

In subsequent analyses we explored which aspects of biological age are marked by ΔmetaboAge. First, we investigated whether ΔmetaboAge correlates with established clinical risk factors for cardio-metabolic disease using phenotypic data available within the BBMRI-NL resource (**Figure 1B**). Meta-analyses across biobanks showed that a positive ΔmetaboAge corresponded with a poor cardio-metabolic health, as represented by higher body mass index (BMI), higher serum levels of C-reactive protein (hsCRP) and not unsurprisingly, higher cholesterol and triglycerides. In addition, use of blood pressure lowering medication, but not lipid lowering medication, is associated with a higher ΔmetaboAge (**Figure 2A**). These associations remained significant when further adjusted for sex and BMI (see **Table S4**).

**FIGURE 2:**
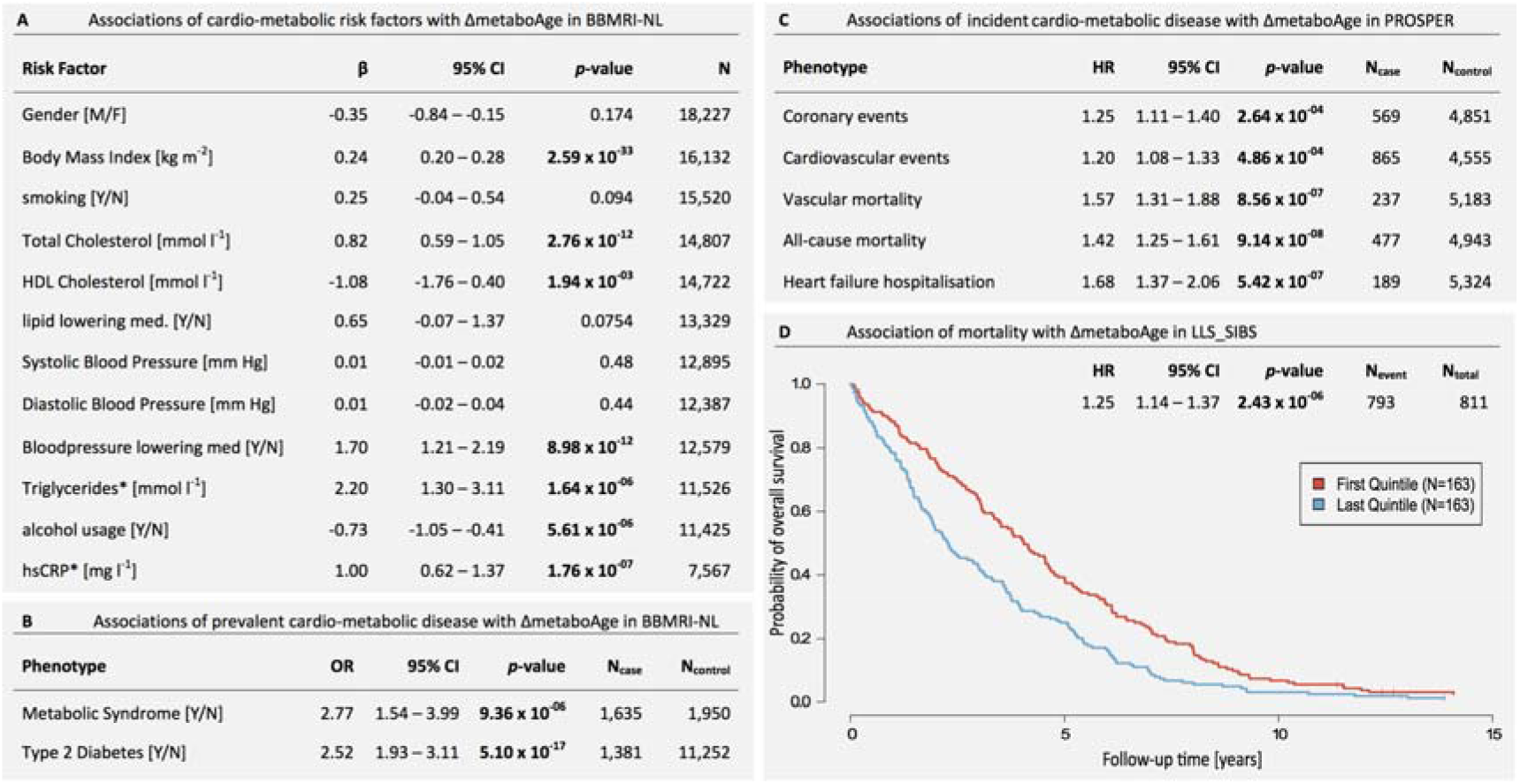
Associations of ΔmetaboAge with (risk factors of) cardio-metabolic disease risk and all-cause mortality. Associations with **A:** Association of cardio-metabolic risk factors with ΔmetaboAge in BBMRI-NL. **B:** Association of prevalent cardio-metabolic disease with ΔmetaboAge in BBMRI-NL. **C:** Association of incident cardio-metabolic disease with ΔmetaboAge in PROSPER. **D:** Association of mortality with ΔmetaboAge in LLS_SIBS adjusted for age and sex. A Kaplan-Meijer curve illustrates the difference in mortality between quintiles with the highest (blue; estimated ≥ 6.9 years older) and lowest (red; estimated ≥ 7.3 years younger) ΔmetaboAge. β’s are reported as increase in ΔmetaboAge per unit of increase in the risk factor (A) or disease status (B). HR’s reported as increased risk per 10-years of ΔmetaboAge. *log-transformed; Cl: confidence interval; med.: medication. P-values are in bold when significant after correction for multiple testing (Bonferroni).

### Associations of metaboAge with current and future cardio-metabolic disease

Next, we investigated whether ΔmetaboAge marks current and future clinical metabolic disease endpoints. Participants with current metabolic syndrome or diabetes mellitus type 2 (T2D) were consistently estimated older as compared to their healthy counterparts of similar age (**Figure 2B**), with T2D remaining significant when also adjusting for sex and BMI (See **Table S5** in **Online Material**). The predictive value of ΔmetaboAge for future cognitive and cardio-metabolic disease was tested in the PROSPER study, a multi-center clinical trial investigating the efficacy of lipid lowering medication for elderly patients (70 – 82 years) at risk of cardiovascular events followed for a median follow-up time of 3.3 years [14] (**Table S2**). While at most marginal correlations were observed between ΔmetaboAge and measures of cognitive decline at baseline (**Table S6**) or during follow-up (**Table S7**), patients with a positive ΔmetaboAge were shown to be at risk of future coronary and cardiovascular events independent of sex, BMI, smoking status, T2D status, anti-hypertensive medication and pravastatin treatment (**Figure 2C**). Using the same model, we found patients with a positive ΔmetaboAge to be at increased risk of heart failure hospitalization and vascular and all-cause mortality (**Figure 2C**).

### Associations of metaboAge with mortality and functionality in the oldest old

Finally we evaluated whether ΔmetaboAge marks biological ageing near the extremes of human life span. We examined participants of the LLS-SIBS [15], aged 89 years and older and followed during a median follow-up time of 12.4 years for all-cause mortality (**Table S2**). At baseline, a positive ΔmetaboAge correlated with lower Instrumental Activities of Daily Living (IADL, *p*=2.0 × 10^−16^), a measure of physical independence. Moreover, a positive ΔmetaboAge also marked nonagenarians at an increased risk of all-cause mortality (**Figure 2D**) during ten years of follow up, even when adjusting for IADL (**Table S8**).

## DISCUSSION

We present a rich resource of ^1^H-NMR serum metabolomics and routine serum measurements encompassing over 25,000 samples, derived from 26 community and hospital based cohorts (download data access request at bbmri.nl/samples-images-data. Using this resource, we have constructed a score reflecting an individual’s biological age, called metaboAge, and demonstrate that the excess of metaboAge over chronological age (ΔmetaboAge) confers an increased risk for future cardiovascular disease, mortality up to the highest ages and functionality among older adults. Lastly, we have made a web-based tool available at metaboage.researchlumc.nl facilitating an easy incorporation of ΔmetaboAge scores in future epidemiological studies.

We evaluated the applicability of ΔmetaboAge as a biomarker for current and future cardio-metabolic health and disease as the same metabolomics platform has previously been successfully employed to predict outcomes for cardiovascular disease [13] and type 2 diabetes [16]. In line with these papers, we observed that higher ΔmetaboAge indicates various aspects of current and future cardio-metabolic health, including significant associations with body mass index (*p* = 2.59 × 10^−33^), C-reactive protein (*p* = 1.76 × 10^−07^), current Type 2 diabetes (*p* = 5.10 × 10^−17^), future cardiovascular events (*p* = 2.64 × 10 ^04^) and vascular mortality (*p* = 8.56 × 10^−07^). Hence, ΔmetaboAge can be readily explored, also in studies lacking cardio-metabolic risk factors or endpoints, as a surrogate marker to capture some aspects of current or future cardio-metabolic health.

Ideally, biomarkers of biological age are broadly applicable and are thus indicative of one or several of the five health domains as defined by Lara *et al.* [17], Whereas we showed that ΔmetaboAge is indicative of classical biomarkers belonging to the physiological (cardiovascular health), immune (hsCRP), and physical capability domain (IADL), we were unable to establish significant correlations with classical biomarkers of the cognitive or endocrine domain. This was either because we lacked the classical biomarkers, as for the endocrine domain, or that ΔmetaboAge did not correlate with the available classical biomarkers, as for the cognitive domain. Of note is that a measure not available to us, general cognitive ability, has recently been reported to associate with several metabolite measurements of this platform in a large epidemiological study [18]. Hence, we expect that future large-scale metabolomics studies using the Nightingale platform, e.g. the UK Biobank, will shed more light on other aspects of biological age indicated by ΔmetaboAge.

While the blood metabolome can be readily assessed using ^1^H-NMR metabolomics at high throughput, high reproducibility and low costs, no ^1^H-NMR metabolomics clock has to date been made available. We have applied the clock paradigm popularized by the work of Horvath *et al.* [8] to derive such a metabolomics-based predictor of age. Similarly, we have shown that our clock associates with various clinical endpoints including mortality. While clock algorithms have become increasingly popular as a means to perform sample stratification, an important limitation of the clock paradigm remains that it is hard to trace back why such scores reflect aspects of current and future disease, let alone for which disease applications a particular score is most suitable. Hence, newly proposed scores inevitably require additional empirical evidence in other epidemiological cohorts to support its added value. To accommodate future research with ΔmetaboAge, we have made a web-based tool available at metaboage.researchlumc.nl.

In summary, we present a rich resource of ^1^H-NMR serum metabolomics and routine serum measurements encompassing over 25,000 samples (download data access request at bbmri.nl/samples-images-data). Moreover, we illustrate the merit of such a resource by presenting ΔmetaboAge, a novel metabolomics-based indicator of biological age capturing aspects of current and future cardio-metabolic health (*p*redictions available at: metaboage.researchlumc.nl).

## METHODS

Methods, including statements of data availability and any associated accession codes and references, are available in the online version of the paper.

## ACKNOWLEDGMENTS

See **Table S9.**

## AUTHOR CONTRIBUTIONS

EBvdA and PES conceived and wrote the manuscript. EBvdA, ST and JJHBW performed the analyses. MJTR verified and supervised the analyses. PES, CMvD en DIB coordinated the BBMRI-NL consortium. MB, JD, HEDS coordinated sample collection and data acquisition. HM, MS, JJHBW, DC en MAS set up the database and catalogue. PES, FWA, EB, PME, JMG, MAI, MK, IM, SPM, RGHHN, MGN, BWJHP, CDAS, CET, GMT, LM’tH, AMJMvdM, PvdH, ICCvdH, CJHvdK, MMJvG WEvS, CW, AZ, AHZ, NS, JWJ, CMvD en DIB made phenotypic and metabolomics measurements available. All authors discussed the results and contributed to the final manuscript.

## ONLINE METHODS

### STUDY SAMPLE

The study sample was selected from the following 26 Dutch biobanks: MRS, VUNTR, CSF, NESDA, FUNCTGENOMICS, LIFELINES, HARM, ERF, HELIUS, CHECK, STEMI_G IPS-III, LLS_PARTOFFS, GARP, CODAM, BIOMARCS, TMS, DCS, TOMAAT, VUMC-ADC, UCORBIO, RAAK, ALPHAOMEGA, TACTICS, ERGO, PROSPER and LLS_SIBS, jointly participating in the Dutch Biobanking and BioMolecular resources and Research infrastructure (BBMRI-NL: http://www.bbmri.nl). Protocols for these studies were approved by the ethics committees at all involved institutes, and informed consent was obtained from all participants. Individuals with missing phenotypic data on age or sex (92) or aged under 18 years (5) were excluded, leaving 23,590 individuals in the current study. A description of the included cohorts is provided in **Table S2** in the **Supplementary Appendix.**

### METABOLOMICS

Metabolite concentrations were measured in EDTA plasma samples using the commercially available high-throughput proton nuclear magnetic resonance (^1^H-NMR) metabolomics platform of *Nightingale Health ltd.* (Helsinki, Finland). In addition to the quantification of routine lipids, lipoproteins, fatty acid composition, amino acids, ketone bodies and gluconeogenesis-related metabolites, this platform also reports on average sizes of lipoprotein particle subclasses VLDL, LDL, and HDL and their respective lipid content ratios. In total, 226 metabolomic variables are reported for measurements in EDTA plasma, described in more detail elsewhere [1], of which 62 were analyzed in the current study (See **Table S3** in the **Supplementary Appendix** for a full list). The selection of 62 metabolomic variables was based on previous publications [2] and omits derived or highly correlating metabolomic variables from the dataset to prevent instability in the age predictor.

### DATA PREPROCESSING

Cohorts reporting on all 62 metabolomic variables were included, i.e. omitting VUNTR (N=3,564) and CODAM (N=150) missing glutamine and acetoacetate respectively. Metabolomic variables measured at low success rates (<98%) or that frequently failed to reach the detection limit (<99%) were excluded (3-hydroxybutyrate, XXL_VLDL_L, XL_VLDL_L, L_VLDL_L, XL_HDL_L and L_HDL_L). Outlier samples were removed by allowing at most 1 missing value per sample (removing 200), 1 zero per sample (removing 64) and no metabolite levels more than 5 standard deviations away from the overall mean per metabolomics variable (removing 442). Obtained data (19,014 samples x 56 metabolomic variables) was scaled and the remaining 467 missing data points (0.046% of the data) were imputed using nipals of the R package pcaMethods. For more details, See **Supplemental Material Doc S2.**

### STATISTICS

#### Constructing the metabolomics-based age predictor

The metabolomics-based age predictor was trained using a linear model:

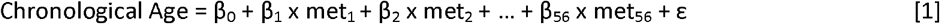

Two evaluation procedures were employed for getting an unbiased estimate of the model performance under two different scenarios. A 5-Fold-Cross-Validation (5FCV) was applied splitting the samples at random in a partition for model training, i.e. ‘TRAIN’ consisting of 80% of the samples, and an independent partition for model evaluation, i.e. ‘TEST’ consisting of 20% of the samples. Samples were drawn while preserving overall age-distribution using the function createFolds of R package caret. Secondly, a Leave-One-Biobank-Out-Validation (LOBOV) was applied by holding out the samples belonging to one particular cohort to create ‘TRAIN’ and evaluating these in the left-out samples, i.e. ‘TEST’.

By lack of a single performance measure reporting faithfully under all conditions, we report two complementary measures, as was previously proposed for the methylation age predictor [3]. First, we report the Pearson correlation between chronological age and predicted age (*r*). Second, we report the median absolute difference between chronological and predicted age (‘median error’). The range and median of both measures are reported across folds or cohorts for respectively the 5CV and LOBOV settings. For more details, See **Supplemental Material Doc S1.**

#### Associations of ΔmetaboAge with other phenotypes and endpoints

To investigate whether age predictions might serve as surrogate biomarkers for biological age and age-related disease, we define Δage as the predicted age minus the actual chronological age, i.e. the model residuals. For direct comparisons, we define ΔmetaboAge as the age-independent part of Δage, obtained by regressing Δage onto chronological age. For associations with phenotypes we prefer to adjust for chronological age directly, e.g.:

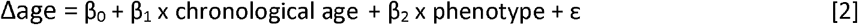

Associations of ΔmetaboAge with phenotypes were performed for each cohort separately and meta-analyzed across cohorts on the estimated coefficients (β_2_) as implemented in the function rma of R package metaphor. While analyses in BBMRI-NL were adjusted for chronological age and optionally sex and BMI, a more extensive model was evaluated in PROSPER to include also the canonical risk factors of cardiovascular disease, i.e. sex, BMI, current smoking status [YES/NO], diabetes mellitus type 2 status [YES/NO], hypertension status [YES/NO] and study covariates, i.e. pravastatin treatment [YES/NO]. Associations with cardio-metabolic phenotypes in BBMRI-NL were summarized using a random effect meta-analysis and β’s were reported as increase in ΔmetaboAge per unit of increase in the risk factor or disease status. The associations with cognitive tests were summarized across the three sub-cohorts of PROSPER using a fixed effect meta-analysis.

A Cox proportional hazards model with age at sampling as the time scale was used to test for the association between ΔmetaboAge and incident endpoints in PROSPER and LLS-SIBS. While we only adjusted for sex and optionally IADL in the LLS_SIBS, we evaluated an extended model in PROSPER representing the aforementioned canonical risk factors for cardiovascular disease and study covariates. HR’s are reported as increased risk per 10-years of ΔmetaboAge.

A linear mixed model was employed to test for the association between differences in time-course trajectories of cognitive tests with ΔmetaboAge:

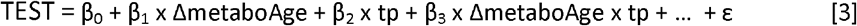

Associations of ΔmetaboAge with cognitive test scores (one of: Stroop, Letter Digit Test (LTD), Picture Learning Test, immediate (PLTi) and delayed (PLTd)) were performed for each sub-cohort of PROSPER separately and fixed-effect meta-analyzed on the estimated coefficients of the interaction term (β_3_) of ΔmetaboAge x tp (timepoint) and were adjusted for the aforementioned canonical risk factors.

#### metaboAge web-based tool

Metabolic age predictions can be obtained at https://bbmri.nl.researchlumc.nl/metaboage. The web-based tool requires the original tab-delimited files as supplied by Nightingale Health and will return metabolic age predictions. Data is transferred to the web-based tool using an encrypted connection (HTTPS), and the data will be stored only the duration of the computation and directly deleted thereafter. In case you would like to contribute your metabolomics data and participate in BBMRI-NL, please contact the corresponding author.

#### Data requests

Please visit bbmri.nl/samples-images-data and fill out and sign the data access request and code of conduct forms to request the data in this manuscript. Applications compliant with ethical and legal legislations will be reviewed by the BBMRI-NL board for overlap with other ongoing projects before access is granted.

## List of Supplemental Material

**Table S1:** BBMRI-NL Consortium banner.

**Table S2:** List of cohort descriptions.

**Table S3:** List of 56 metabolomic variables used to train the model.

**Table S4:** Associations with cardio-metabolic risk factors with ΔmetaboAge in BBMRI-NL adjusted for sex and BMI.

**Table S5:** Associations of T2D status with ΔmetaboAge in BBMRI-NL adjusted for sex and BMI.

**Table S6:** Associations of cognitive measures with ΔmetaboAge in PROSPER adjusted for sex, BMI, current smoking status [YES/NO], prevalence of T2D [YES/NO] and hypertension status [YES/NO] and pravastatin treatment [YES/NO].

**Table S7:** Associations of change in cognitive measures with ΔmetaboAge in PROSPER adjusted for sex, BMI, current smoking status [YES/NO], prevalence of T2D [YES/NO] and hypertension status [YES/NO] and pravastatin treatment [YES/NO].

**Table S8:** Associations of mortality with ΔmetaboAge in LLS_SIBS, adjusted for sex and IADL.

**Table S9:** Acknowledgements.

**Doc S1:** Model training.

**Doc S2:** Quality control.

