## Supplemental Table S1 for "Predicting biological age based on the BBMRI-NL ^1^H-NMR metabolomics repository"

### Table S1 BBMRI-NL

#### BBMRI-NL Consortium

**Analysis Group:** E.B. van den Akker^1-3^, J.J.H. Barkey Wolf^1^, S. Trompet^4,5^, M.J.T. Reinders^2,3^

**Cohort collection and sample management group:**

M. Beekman^1^, H.E.D. Suchiman^1^, J. Deelen^1,6^**,** N. Amin^7^, J.W. Beulens^8,9^, J.A. van der Bom^10-13^, N. Bomer^14^, A. Demirkan^7^, J.A. van Hilten^15^, J.M.T.A. Meessen^16^, R. Pool^17^, M.H. Moed^1^, J. Fu^18,19^, G.L.J. Onderwater^20^, F. Rutters^8^, C. So-Osman^10^, W.M. van der Flier^8,21^, A.A.W.A. van der Heijden^22^, A. van der Spek^7^**,** F.W. Asselbergs^23^, E. Boersma^24^, P.M. Elders^25,26^, J.M. Geleijnse^27^, M.A. Ikram^7,29,30^, M. Kloppenburg^13,31^, I. Meulenbelt^1^, S.P. Mooijaart^4^, R.G.H.H. Nelissen^32^, M.G. Netea^33,34^, B.W.J.H. Penninx^26,35^, C.D.A. Stehouwer^36,37^, C.E. Teunissen^38^, G.M. Terwindt^20^, L.M. ‘t Hart^1,8,26,39,40^, A.M.J.M. van den Maagdenberg^41^, P. van der Harst^14^, I.C.C. van der Horst^42^, C.J.H. van der Kallen^36,37^, M.M.J. van Greevenbroek^36,37^, W.E. van Spil^43^, C. Wijmenga^18^, A.H. Zwinderman^44^, A. Zhernikova^18^, J.W. Jukema^5^, N. Sattar^45^

**Database & Catalogue:** J.J.H. Barkey Wolf^1^, M. Beekman^1^, D. Cats^1^, H. Mei^1,46^, M. Slofstra^18^, M. Swertz^18^,

**Steering committee:** D.I. Boomsma^26,47^, C.M. van Duijn^7^, P.E. Slagboom^1^

##### Affiliations

1. Department of Molecular Epidemiology, Leiden University Medical Center, Leiden, The Netherlands
2. Leiden Computational Biology Center, Leiden University Medical Center, Leiden, The Netherlands
3. Department of Pattern Recognition and Bioinformatics, Delft University of Technology, Delft, The Netherlands
4. Department of Internal Medicine, Division of Gerontology and Geriatrics, Leiden University Medical Centre, Leiden, The Netherlands
5. Department of Cardiology, Leiden University Medical Center, Leiden, The Netherlands
6. Max Planck Institute for Biology of Ageing, Cologne, Germany
7. Department of Epidemiology, Erasmus MC University Medical Center, Rotterdam, The Netherlands
8. Department of Epidemiology and Biostatistics, Amsterdam University Medical Center, Amsterdam, the Netherlands
9. Julius Center for Health Sciences and Primary Care, University Medical Center Utrecht, Utrecht, The Netherlands.
10. Centre for Clinical Transfusion Research, Sanquin Research, Leiden, The Netherlands.
11. Jon J van Rood Centre for Clinical Transfusion Research, Leiden University Medical Centre, Leiden, The Netherlands.
12. TIAS, Tilburg University, Tilburg, The Netherlands.
13. Department of Clinical Epidemiology, Leiden University Medical Centre, Leiden, The Netherlands.
14. Department of Cardiology, University Medical Center Groningen, University of Groningen, Groningen, the Netherlands.
15. Center for Clinical Transfusion Research, Sanquin Research, Leiden, the Netherlands.
16. Department of Orthopedics, Leiden University Medical Centre, Leiden, The Netherlands.
17. Department of Biological Psychology, Vrije Universiteit, Amsterdam, the Netherlands.
18. Department of Genetics, University Medical Center Groningen, University of Groningen, Groningen, The Netherlands
19. Department of Pediatrics, University Medical Center Groningen, University of Groningen, Groningen, The Netherlands
20. Department of Neurology, Leiden University Medical Center, Leiden, the Netherlands.
21. Department of Neurology and Alzheimer Center, Neuroscience Campus Amsterdam, VU University Medical Center, Amsterdam, The Netherlands.
22. Department of General Practice, The EMGO Institute for Health and Care Research, VU University Medical Center, Amsterdam, The Netherlands.
23. Department of Cardiology, Division Heart and Lungs, University Medical Center Utrecht, Utrecht, The Netherlands Julius Center for Health Sciences and Primary Care, University Medical Center Utrecht, Utrecht, The Netherlands.
24. Thorax centre, Erasmus Medical Centre, Rotterdam, the Netherlands
25. Department of General Practice and Elderly Care Medicine, VU University Medical Center, Amsterdam, The Netherlands
26. Amsterdam Public Health research institute, VU University Medical Center, Amsterdam, The Netherlands
27. Division of Human Nutrition and Health, Wageningen University, Wageningen, The Netherlands
28. Department of Epidemiology, Erasmus University Medical Center Rotterdam, Rotterdam, The Netherlands
29. Department of Radiology, Erasmus University Medical Center Rotterdam, Rotterdam, The Netherlands
30. Department of Neurology, Erasmus University Medical Center Rotterdam, Rotterdam, The Netherlands
31. Department of Rheumatology, Leiden University Medical Center, The Netherlands
32. Department of Orthopaedics, Leiden University Medical Center, Leiden, The Netherlands
33. Department of Internal Medicine, Radboud Center for Infectious Diseases, Radboud University Medical Center, Nijmegen, Netherlands
34. Department for Genomics & Immunoregulation, Life and Medical Sciences Institute (LIMES), University of Bonn, Bonn, Germany
35. Department of Psychiatry, VU University Medical Center, Amsterdam, The Netherlands.
36. Department of Internal Medicine, Maastricht University Medical Center (MUMC+), Maastricht, the Netherlands
37. School for Cardiovascular Diseases (CARIM), Maastricht University, Maastricht, the Netherlands
38. Neurochemistry Laboratory, Clinical Chemistry Department, Amsterdam University Medical Center, Amsterdam Neuroscience, the Netherlands
39. Department of Cell and Chemical Biology, Leiden University Medical Center, Leiden, the Netherlands
40. Department of General practice, Amsterdam University Medical Center, Amsterdam, the Netherlands
41. Department of Human Genetics, Leiden University Medical Center, Leiden, The Netherlands
42. Department of Critical Care, University Medical Center Groningen, Groningen, The Netherlands.
43. UMC Utrecht, Department of Rheumatology & Clinical Immunology, Utrecht, The Netherlands
44. Department of Clinical Epidemiology, Biostatistics, and Bioinformatics, Academic Medical Centre, University of Amsterdam, Amsterdam, the Netherlands
45. Institute of Cardiovascular and Medical Sciences, Cardiovascular Research Centre, University of Glasgow, Glasgow, UK
46. Sequencing Analysis Support Core, Leiden University Medical Center, Leiden, The Netherlands
47. Netherlands Twin Register, Department of Biological Psychology, Vrije Universiteit, Amsterdam, The Netherlands
