## Supplemental Table S2 for "Predicting biological age based on the BBMRI-NL ^1^H-NMR metabolomics repository"

**Table S2:** Cohort Descriptions

**Alpha Omega**

The Alpha Omega Cohort consists of 4,837 Dutch men and women aged 60-80 years with a clinically diagnosed myocardial infarction <10 years before study enrollment. Baseline examinations and blood sampling took place between 2002 and 2006, after which the cohort has been followed up for cause-specific mortality. During the first 3 years of follow-up, patients participated in an intervention study with omega-3 fatty acids (Alpha Omega Trial) [1,2]. For the present analysis, a random sample of 600 patients was selected, who were followed up through Statistics Netherlands (CBS) until the 1^st^ of January 2014. Metabolites were successfully quantified in EDTA plasma samples of 568 patients. The Alpha Omega Cohort is registered with clinicaltrials.gov (Identifier: NCT03192410).

1. Geleijnse JM, Giltay EJ, Schouten EG, et al. Effect of low doses of n-3 fatty acids on cardiovascular diseases in 4,837 post-myocardial infarction patients: design and baseline characteristics of the Alpha Omega Trial. Am Heart J 2010;159:539-46 e2.

2. Kromhout D, Giltay EJ, Geleijnse JM, Alpha Omega Trial G. n-3 fatty acids and cardiovascular events after myocardial infarction. N Engl J Med 2010;363:2015-26.

**Amsterdam Dementia Cohort**

The Amsterdam Dementia Cohort is an ongoing study including patients who visit the memory clinic of the Alzheimer center of the VU University Medical Center [1]. At baseline, all subjects receive a diagnostic assessment including medical history taking, physical and neurological examination, neuropsychological investigation, standard laboratory tests of blood and cerebrospinal fluid, electroencephalogram and brain magnetic resonance imaging. Clinical diagnosis is made in consensus based, multidisciplinary meetings. For the present metabolomics analysis, 1,473 plasma samples were selected based on available volume in our biobank. In this cohort 45% is female and mean±SE age of 64±9. Date of enrollment was between 2001-2014.

1. van der Flier, W.M., et al., Optimizing patient care and research: the Amsterdam Dementia Cohort. J Alzheimers Dis, 2014. 41(1): p. 313-27.

**BIOMARCS**

The BIOMarker study to identify the Acute risk of a Coronary Syndrome (BIOMArCS) was designed to study the relation between temporal changes in cardiovascular biomarkers and ischemic cardiovascular events in patients discharged after acute coronary syndrome (ACS) admission.[1] 844 ACS patients were enrolled in 18 hospitals in The Netherlands. Venipuncture was scheduled at 19 regular intervals during a year. 45 Patients (cases) reached the study endpoint of repeat ACS within one year. BIOMArCS was approved by the institutional review committees of the participating hospitals. All patients gave informed consent.

1. Rohit M Oemrawsingh et al. , Cohort profile of BIOMArCS: the BIOMarker study to identify the Acute risk of a Coronary Syndrome-a prospective multicentre biomarker study conducted in the Netherlands; BMJ Open. 2016 Dec 23;6(12):e012929. doi: 10.1136/bmjopen-2016-012929.

**LUMINA**

The Leiden University MIgraine Neuro-Analysis (LUMINA) cohort currently consists of over 6,700 male and female participants aged 18-88 years. LUMINA participants are recruited through a dedicated, nationwide website (<http://www.lumc.nl/org/hoofdpijn-onderzoek/onderzoek>) inviting Dutch migraine patients and non-migraine controls to participate in migraine research. Participants recruited through the website were asked to complete a screening questionnaire, previously validated to diagnose migraine [1]. The questionnaire was validated by a semi-structured telephone interview, in accordance with the International Classification of Headache Disorders (ICDH-3β)[2]. In addition, patients attending the Leiden University Medical Centre (LUMC) dedicated headache clinic were invited to participate. Migraine patients recruited through the headache clinic were diagnosed with migraine by a neurologist specialized in headache. For the present metabolomics analysis, a sample of migraine patients and non-migraine controls was selected, who were participating in several LUMINA sub cohorts (CSF, MRS, and CHARM cohorts) between 2008-2014. Migraine diagnosis was confirmed by the study-physician at the day of blood draw resulting in the collection of 564 blood (EDTA/Serum) samples from 432 individual participants used for metabolite quantification.

1. van Oosterhout WPJ, Weller CM, Stam AH, et al. Validation of the web-based LUMINA questionnaire for recruiting large cohorts of migraineurs. Cephalalgia. 2011;31:1359–1367. doi: 10.1177/0333102411418846.

2. Headache Classification Committee of the International Headache Society (IHS). The International Classification of Headache Disorders, 3rd edition (beta version). Cephalalgia. 2013;33:629–808. doi: 10.1177/0333102413485658.

**CHECK**

CHECK (Cohort Hip & Cohort Knee) is a prospective, 10-year follow-up, observational cohort study of 1002 people aged between 45 and 65 years at the time of inclusion, with pain in their knee(s) and/or hip(s), who had never or not longer than 6 months ago consulted a physician for these complaints [1]. Blood samples were taken non-fasted. Hip and knee radiographs were obtained multiple times throughout follow-up and scored pairwise according to the Kellgren & Lawrence (KL) scoring system. When scored pairwise, these people did not have obvious radiographic knee or hip OA at baseline (i.e. KL0 or 1).

1. Wesseling J, Dekker J, van den Berg WB, Bierma-Zeinstra SM, Boers M, Cats HA, et al. CHECK (Cohort Hip and Cohort Knee): similarities and differences with the Osteoarthritis Initiative. Ann Rheum Dis 2009;68:1413-9.

**CODAM**

The Cohort on Diabetes and Atherosclerosis Maastricht [1] (CODAM) is a prospective, observational study that consists of 574 individuals who were selected from a larger population-based cohort [2]. Inclusion of participants into CODAM was based on a moderately increased risk to develop cardiometabolic diseases, such as type 2 diabetes and/or cardiovascular disease. Participants were included if they were of Caucasian descent and over 40 yrs of age and additionally met at least one of the following criteria: increased BMI (>25 kg/m^2^), a positive family history of type 2 diabetes, a history of gestational diabetes and/or glycosuria, or use of anti-hypertensive medication. The CODAM baseline measurements were done between 2000 and 2002 (n=352 men, 222 women, age 59.6±7.0 yrs). After a median of 7.0 years (interquartile range 6.9–7.1), 495 participants were included in the follow-up measurements [3]. Metabolites were obtained fasting EDTA plasma samples of all participants at baseline and at follow-up in those participants who had type 2 diabetes at baseline (n=110).

1. van Greevenbroek, M. M. J. et al. The cross-sectional association between insulin resistance and circulating complement C3 is partly explained by plasma alanine aminotransferase, independent of central obesity and general inflammation (the CODAM study). Eur. J. Clin. Invest. 41, 372–379 (2011). doi: 10.1111/j.1365-2362.2010.02418.x

2. Van Dam, R. M., Boer, J. M., Feskens, E. J. M. & Seidell, J. C. Parental history of diabetes modifies the association between abdominal adiposity and hyperglycemia. Diabetes Care 24, 1454–1459 (2001).

3. Wlazlo N, van Greevenbroek MM, Ferreira I, Feskens EJ, van der Kallen CJ, Schalkwijk CG, Bravenboer B, Stehouwer CD. Complement factor 3 is associated with insulin resistance and with incident type 2 diabetes over a 7-year follow-up period: the CODAM Study. Diabetes Care37, 1900-1909 (2014). doi: 10.2337/dc13-2804

**The Maastricht Study (TMS)**

The Maastricht Study [1] is an observational prospective population-based cohort study enriched with T2DM individuals. Eligible for participation were individuals aged between 40 and 75 years and living in the southern part of the Netherlands (municipalities Maastricht, Margraten-Eijsden, Meersen and Valkenburg; Maastricht and Heuvelland in the province of Limburg).

1. Schram MT1 et al. The Maastricht Study: an extensive phenotyping study on determinants of type 2 diabetes, its complications and its comorbidities. Eur J Epidemiol. 2014 Jun;29(6):439-51. doi: 10.1007/s10654-014-9889-0. Epub 2014 Apr 23.

**The Hoorn Diabetes Care System cohort study (DCS)**

The DCS cohort currently consists of approximately 13.000people with type 2 diabetes. The DCS provides routine diabetes care to people with type 2 diabetes living in the West-Friesland region of the Netherlands [1]. People treated by the DCS visit the DCS research center annually, during which blood is drawn in the fasting state for routine biochemistry. Furthermore, all participants get a full medical exam, advice about their health and treatment and receive education on their disease during their annual visits to the DCS research center. In addition, patients are invited to join our research and biobanking studies (n=5,000+). From the DCS biobank we included for this study a random cross-sectional sample for which a baseline plasma sample and yearly follow-up data were available (n=750). For case-control analyses this sample was supplemented with individuals selected for the inability to reach the glycaemic target (HbA1c>53 mmol/mol) and/or suffering from diabetic complications (n=245). Samples were collected in 2008/2009 and stored and -80 degrees Celsius until analysis. Metabolites were successfully quantified in fasting EDTA plasma samples from 995 individuals.

[1] van der Heijden AA, Rauh SP, Dekker JM, Beulens JW, Elders P, 't Hart LM, Rutters F, van Leeuwen N, Nijpels G: The Hoorn Diabetes Care System (DCS) cohort. A prospective cohort of persons with type 2 diabetes treated in primary care in the Netherlands. BMJ Open 2017;7:e015599

**Website**: [www.hoornstudies.com](http://www.hoornstudies.com)

**Erasmus Rucphen Family study (ERF)**

The Erasmus Rucphen Family is a family-based study that includes inhabitants of a genetically isolated community in the South-West of the Netherlands [1]. The goal of the study is to identify the risk factors in the development of complex disorders. Study population includes approximately 3,000 individuals who are living descendants of a limited number of founders living in the 19^th^ century [1]. All data were collected between 2002 and 2005. All participants gave informed consent, and the Medical Ethics Committee of the Erasmus University Medical Centre approved the study.

Metabolomics measurements were quantified from fasted EDTA plasma samples using Nightingale Health platform. Metabolomics measurements were available for 1,402 participants from the ERF.

1. Pardo, Luba M., et al. "The effect of genetic drift in a young genetically isolated population." Annals of human genetics69.3 (2005): 288-295.

**Rotterdam Study (RS)**

The Rotterdam Study is a prospective, population-based cohort study among individuals living in the well-defined Ommoord district in the city of Rotterdam in The Netherlands [1]. The aim of the study is to determine the occurrence of cardiovascular, neurological, ophthalmic, endocrine, hepatic, respiratory, locomotor, dermatological, otolaryngological, and psychiatric diseases in elderly people. The cohort was initially defined in 1990 among approximately 7,983 persons, aged 55 years and older, who underwent a home interview and extensive physical examination at the baseline and during follow-up visits every 3-4 years (RS-I)[1]. Cohort was extended in 2000/2001 (RS-II, 3,011 individuals aged 55 years and older) and 2006/2008 (RS-III, 3,932 subjects, aged 45 and older). As of 2008, Rotterdam Study comprised 14,926 subjects. Written informed consent was obtained from all participants and the Medical Ethics Committee of the Erasmus Medical Center, Rotterdam, approved the study. Metabolomics measurements were quantified in fasted EDTA plasma samples using Nightingale Health platform. Metabolomics measurements were available for 2,986 participants from RS-I, 591 participants from RS-II (n=591) and 1,787 participants from RS-III.

1. Ikram, M. Arfan, et al. "The Rotterdam Study: 2018 update on objectives, design and main results." *European Journal of Epidemiology* 32.9 (2017): 807-850.

**FUNCTGENOMICS**

The 500 Functional Genomics (500FG [1]) project consists of 534 adult healthy volunteers sampled between July 2013 and December 2014. Inclusion criteria were >18 years of age and Western European descent. Exclusion criteria were pregnancy/breastfeeding, chronic or acute disease at the time of assessment, and use of chronic or acute medication during the last month before the study. After visiting the hospital to donate blood, volunteers received an extensive online questionnaire about lifestyle, diet, and disease history. Upon analyzing the questionnaire data we excluded 45 volunteers as they were under medication, non-European descent, or had kidney disease or diabetes mellitus.

1. Schirmer et al. Linking the Human Gut Microbiome to Inflammatory Cytokine Production Capacity. Cell. 2016 Nov 3;167(4):1125-1136.e8. doi: 10.1016/j.cell.2016.10.020.

**GARP**

The GARP cohort (N=217) consists of patients with advanced radiographic OA at two or more joint sites of hand, spine, knee or hip. Follow-up was performed at 5 years, at which radiographs for hip, knee and hand were scored pairwise using the OARSI Atlas and the KL scoring system. Matched to the GARP study, a normal reference control group (NORREF) was collected using the same protocol and included in this study as controls [1-3]. Blood was collected non-fasted.

1. Riyazi N, Meulenbelt I, Kroon HM, Ronday KH, Hellio le Graverand MP, Rosendaal FR, et al. Evidence for familial aggregation of hand, hip, and spine but not knee osteoarthritis in siblings with multiple joint involvement: the GARP study. Ann Rheum Dis 2005;64:438-43.

2. Meulenbelt I, Kloppenburg M, Kroon HM, Houwing-Duistermaat JJ, Garnero P, Hellio-Le Graverand MP, et al. Clusters of biochemical markers are associated with radiographic subtypes of osteoarthritis (OA) in subject with familial OA at multiple sites. The GARP study. Osteoarthritis Cartilage 2007;15:379-85.

3. Bijsterbosch J, Meulenbelt I, Watt I, Rosendaal FR, Huizinga TW, Kloppenburg M. Clustering of hand osteoarthritis progression and its relationship to progression of osteoarthritis at the knee. Ann Rheum Dis 2014;73:567-72.

**HELIUS study**

The HELIUS study is a prospective cohort study among the largest ethnic groups living in Amsterdam, the Netherlands. The aim of the HELIUS study is to investigate the causes of (the unequal burden of) diseases across ethnic groups, focusing on three disease categories: cardiovascular diseases, mental health and infectious diseases [1]. Between 2011-2015, a total 24,789 participants (men and women aged 18-70 years) were included at baseline. Similar-sized samples of individuals of Dutch, African Surinamese, South-Asian Surinamese, Ghanaian, Turkish and Moroccan origin were included. Participants filled in an extensive questionnaire and underwent a physical examination that included the collection of biological samples (biobank). Follow-up data is obtained by linkages with existing registries (e.g. hospital data, insurance data) and will be obtained by repeated measurements [2]. Metabolites were quantified in EDTA plasma samples of 500 African origin participants with (pre)diabetes (235 African Surinamese, 265 Ghanaian participants).

[1] K Stronks, MB Snijder, RJ Peters, M Prins, AH Schene, AH Zwinderman. Unravelling the impact of ethnicity on health in Europe: the HELIUS study. BMC Public Health 2013;13:402.

[2] MB Snijder, H Galenkamp, M Prins, EM Derks, RJ Peters, AH Zwinderman, K Stronks. Cohort Profile: the Healthy Life in an Urban Setting (HELIUS) study. BMJ Open (in press).

**Website:** [www.heliusstudie.nl](http://www.heliusstudie.nl)

**LIFELINES-DEEP**

The LifeLines-DEEP cohort is a subset of the Dutch general population cohort LifeLines. Both LifeLines and LifeLines DEEP have been previously described [1-3]. In summary, LifeLines is a three-generation observational follow-up study, which was set up to investigate universal risk factors and their modifiers for multifactorial diseases. Since 2006, approximately 167,000 individuals from the general population residing in the three northern provinces of the Netherlands participate in the study. All participants will be followed-up prospectively for at least 30 years. Participants regularly undergo physical examinations and fill in extensive questionnaires. In addition, blood and urine samples are collected. Each participant is asked to fill in health, lifestyle, and quality-of-life questionnaires every 1.5 years, whereas each participant is invited for a follow-up visit to a Lifelines clinic every 5 years [1,2]. LifeLines-DEEP comprises 1,539 participants (636 males and 903 females, age range 18–84 years). This study was set up for the more detailed phenotyping and omics profiling. For analysis of the genome, epigenome, transcriptome, microbiome, metabolome and other biological levels, additional biomaterials were collected, including additional blood, exhaled air and fecal samples, as well as responses to gastrointestinal health [3].

The current metabolomics study included the baseline information and plasma samples of LifeLines-DEEP participants. EDTA plasma samples were collected after overnight fasting. Peripheral blood samples were drawn by venipuncture from the median cubital vein and subsequently placed at 4°C. Transport of the samples from the research site to the LifeLines laboratory in Groningen was under tightly controlled and continuously monitored conditions. At the LifeLines site, plasma was prepared and aliquoted and stored at -80°C. The samples underwent two freeze-thaw cycles prior to shipment to Brainshake for metabolome analysis [1]. With some sample drop-off, this study eventually included 1,440 LifeLines-DEEP participants. The LifeLines DEEP study was approved by the institutional ethics review board of University Medical Center Groningen (ref. M12.113965).

1. Scholtens, S. et al. Cohort Profile: LifeLines, a three-generation cohort study and biobank. Int. J. Epidemiol. 44, 1172–1180 (2015).

2. Stolk, R. P. et al. Universal risk factors for multifactorial diseases: LifeLines: A three-generation population-based study. Eur. J. Epidemiol. 23, 67–74 (2008).

3. Tigchelaar E. F. et al., Cohort profile: LifeLines DEEP, a prospective, general population cohort study in the northern Netherlands: study design and baseline characteristics. BMJ Open 5, e006772 (2015).

**Website:** <https://www.lifelines.nl/>

**Leiden Longevity Study (LLS)**

The Leiden Longevity Study (LLS) consists of 421 long-lived families of European descent. Families were included if at least two long-lived siblings were alive and fulfilled the age criterion of 89 years or older for males and 91 years or older for females, representing <0·5% of the Dutch population in 2001 [1]. In total, 944 long-lived proband siblings (mean age = 94 years, range = 89-104), 1671 offspring (mean age = 61 years, range = 39-81) and 744 spouses thereof (mean age = 60 years, range = 36-79) were included. Registry-based follow-up until the 27^th^ of October 2016 was available for all participants. Metabolites were successfully quantified in 843 nonagenarians [LLS_SIBS], 1157 of their offspring and 684 controls (LLS_PAROFF) using non-fasted EDTA plasma samples.

1. M. Schoenmaker et al. Evidence of genetic enrichment for exceptional survival using a family approach: the Leiden Longevity Study. Eur J Hum Genet. 2006 Jan;14(1):79-84.

**NESDA**

The Netherlands Study of Depression and Anxiety (NESDA) is an ongoing observational longitudinal cohort study on the long term course and consequences of depressive and anxiety disorders [1]. Between September 2004 and February 2007, 2,981 participants (1,979 females, 1,002 males) aged 18 through 65 years were included. They were recruited through different settings (community, primary care and specialized mental health clinics) in order to obtain a representative sample of persons with depressive and/or anxiety disorders (in lifetime, n=2,329) and without depressive and/or anxiety disorders (n=652). At baseline, participants completed the 4-hour baseline assessment, which included a face-to-face interview, written questionnaires, and biological measurements. Follow-up visits to the research center have now been finished 2, 4, 6 and 9 years after baseline, with response rates of n=2,596, n=2,402, n=2,256 and n=2,069, respectively. The research protocol was approved by the Ethical Committee of the participating centers, and all participants provided written informed consent. During the baseline interview, EDTA plasma samples were collected and stored in aliquots at -85°C until further analysis. Participants were instructed to have an overnight fast before blood collection. Metabolites in these blood samples were analyzed in 2 batches (April and December 2014, respectively) by Brainshake Ltd./Nightingale Health, Helsinki, Finland.

[1] Penninx BWJH, Beekman ATF, Smit JH, Zitman FG, Nolen WA, Spinhoven P, et al. The

Netherlands Study of Depression and Anxiety (NESDA): rationale, objectives and methods. Int J Methods Psychiatr Res 2008;17:121–40. doi: 10.1002/mpr.256.

**Website:** [**http://www.nesda.nl**](http://www.nesda.nl)

**PROspective Study of Pravastatin in the Elderly at Risk (PROSPER)**

The PROspective Study of Pravastatin in the Elderly at Risk (PROSPER) trial design has been published [1,2]. In brief, 5,804 elderly adults (70-82 years old) were enrolled. This was a double-blind, randomised placebo controlled trial investigating the benefit of pravastatin (40 mg/day) in elderly individuals at risk of CVD. Participants were identified in the primary care setting from 3 centres: Glasgow, Scotland; Cork, Ireland or Leiden, the Netherlands. All participants had high-normal to high cholesterol (4·0-9·0 mmol/L) at baseline. Additionally 50% of patients had evidence of vascular disease (physician diagnosed stable angina, stroke, transient ischaemic attack (TIA) or myocardial infarction (MI)) and the remaining 50% of patients had high risk of vascular disease as they had either hypertension, diabetes or were smokers. The primary outcome measure of PROSPER was a composite CVD outcome. In the current study the endpoint of interest was all-cause mortality. Patients were recruited between December 1997 and May 1999 and the mean follow-up period was 3·2 years. Fasting venous blood samples were collected at baseline and at 3-month intervals and biobanked at -80°C. For the present study previously unthawed 6-month post-randomisation samples were used, employing the study as a cohort study and adjusting for randomised treatment in the analyses. Metabolites were successfully quantified in 5,329 individuals.

1. Shepherd J, Blauw GJ, Murphy MB, Cobbe SM, Bollen EL, Buckley BM, et al. The design of a prospective study of Pravastatin in the Elderly at Risk (PROSPER). PROSPER Study Group. PROspective Study of Pravastatin in the Elderly at Risk. Am J Cardiol 1999 Nov 15;84(10):1192-7. PMID: 10569329

2. Shepherd J, Blauw GJ, Murphy MB, Bollen EL, Buckley BM, Cobbe SM, et al. Pravastatin in elderly individuals at risk of vascular disease (PROSPER): a randomised controlled trial. Lancet 2002 Nov 23;360(9346):1623-30. PMID:12457784

**LUMC Arthroplasty studies:**

The LUMC arthroplasty studies (N=462) consist of participants of the RAAK, TacTics (NTR309) and TOMaat (NTR303) studies [1, 2]. These cross-sectional studies included OA patients who received THA or TKA. Since all participants underwent THA/TKA, all patients are considered as end-stage OA and included in the cross-sectional OA prevalence analysis. Blood samples were collected during surgery while patients were fasted.

[1] Ramos YF, den Hollander W, Bovee JV, Bomer N, van der Breggen R, Lakenberg N, et al. Genes involved in the osteoarthritis process identified through genome wide expression analysis in articular cartilage; the RAAK study. PLoS One 2014;9:e103056.

[2] So-Osman C, Nelissen RG, Koopman-van Gemert AW, Kluyver E, Poll RG, Onstenk R, et al. Patient blood management in elective total hip- and knee-replacement surgery (Part 1): a randomized controlled trial on erythropoietin and blood salvage as transfusion alternatives using a restrictive transfusion policy in erythropoietin-eligible patients. Anesthesiology 2014;120:839-51.

**STEMI-GIPS-III**

The Glycometabolic Intervention as Adjunct to Primary Coronary Intervention in ST Elevation Myocardial Infarction (GIPS-III) study is a placebo-controlled randomized clinical trial to evaluate the effect of metformin therapy on left ventricular function in 380 non-diabetic ST-elevated myocardial infarction (STEMI) patients aged 23-90. Patients were included between 2011 and 2013. Blood samples were collected at several time points after inclusion. During the 4-month treatment period, patients received metformin or placebo. 4 months after randomization, left ventricular ejection fraction (LVEF) was measured by cardiac MRI. In addition, patients were followed up for major adverse cardiac events (death, recurrent MI, target lesion revascularization), stroke, non-elective hospitalizations for chest pain or heart failure, all recurrent coronary interventions and internal cardiac defibrillator implantations. Metabolic profiling was assessed in EDTA plasma samples collected at baseline (hospital admission), 24 h post-MI and 4 months post-MI. GIPS-III is registered with clinicaltrials.gov (ID: NCT01217307)

1. Eppinga RN, Kofink D, Dullaart RP, Dalmeijer GW, Lipsic E, van Veldhuisen DJ, van der Horst IC, Asselbergs FW, van der Harst P. Effect of Metformin on Metabolites and Relation With Myocardial Infarct Size and Left Ventricular Ejection Fraction After Myocardial Infarction. Circ Cardiovasc Genet. 2017 Feb;10(1).

2. Eppinga RN, Hartman MH, van Veldhuisen DJ, Lexis CP, Connelly MA, Lipsic E, van der Horst IC, van der Harst P, Dullaart RP. Effect of Metformin Treatment on Lipoprotein Subfractions in Non-Diabetic Patients with Acute Myocardial Infarction: A Glycometabolic Intervention as Adjunct to Primary Coronary Intervention in ST Elevation Myocardial Infarction (GIPS-III) Trial. PLoS One. 2016 Jan 25;11(1):e0145719.

3. Lexis CP, van der Horst-Schrivers AN, Lipsic E, Valente MA, Muller Kobold AC, de Boer RA, van Veldhuisen DJ, van der Harst P, van der Horst IC. The effect of metformin on cardiovascular risk profile in patients without diabetes presenting with acute myocardial infarction: data from the Glycometabolic Intervention as adjunct to Primary Coronary Intervention in ST Elevation Myocardial Infarction (GIPS-III) trial. BMJ Open Diabetes Res Care. 2015 Dec 11;3(1):e000090.

4. Lexis CP, van der Horst IC, Lipsic E, Wieringa WG, de Boer RA, van den Heuvel AF, van der Werf HW, Schurer RA, Pundziute G, Tan ES, Nieuwland W, Willemsen HM, Dorhout B, Molmans BH, van der Horst-Schrivers AN, Wolffenbuttel BH, ter Horst GJ, van Rossum AC, Tijssen JG, Hillege HL, de Smet BJ, van der Harst P, van Veldhuisen DJ; GIPS-III Investigators. Effect of metformin on left ventricular function after acute myocardial infarction in patients without diabetes: the GIPS-III randomized clinical trial. JAMA. 2014 Apr 16;311(15):1526-35.

5. Lexis CP, van der Horst IC, Lipsic E, van der Harst P, van der Horst-Schrivers AN, Wolffenbuttel BH, de Boer RA, van Rossum AC, van Veldhuisen DJ, de Smet BJ; GIPS-III Investigators. Metformin in non-diabetic patients presenting with ST elevation myocardial infarction: rationale and design of the glycometabolic intervention as adjunct to primary percutaneous intervention in ST elevation myocardial infarction (GIPS)-III trial. Cardiovasc Drugs Ther.

**UCORBIO**

The Utrecht Coronary Biobank Study (UCORBIO) enrolled 2,591 patients aged 18-93 who underwent coronary angiography for any indication at the UMC Utrecht. Baseline assessment and blood sampling took place between 2011 and 2014. Patients were followed up for the occurrence of major adverse cardiovascular events (stroke, myocardial infarction, coronary revascularization, death). During the follow-up period (maximum: 3 years), patients completed questionnaires every year to obtain information on hospital admissions. General practitioners and hospitals were contacted to confirm reported cardiovascular events. EDTA samples of 1,198 patients were selected for metabolic profiling. UCORBIO is registered with clinicaltrials.gov (ID: NCT02304744).

1. Gijsberts CM, Santema BT, Asselbergs FW, de Kleijn DP, Voskuil M, Agostoni P, Cramer MJ, Vaartjes I, Hoefer IE, Pasterkamp G, den Ruijter HM. Women Undergoing Coronary Angiography for Myocardial Infarction or Who Present With Multivessel Disease Have a Poorer Prognosis Than Men. Angiology. 2016 Jul;67(6):571-81.

2. Gijsberts CM, Agostoni P, Hoefer IE, Asselbergs FW, Pasterkamp G, Nathoe H, Appelman YE, de Kleijn DP, den Ruijter HM. Gender differences in health-related quality of life in patients undergoing coronary angiography. Open Heart. 2015 Aug 27;2(1):e000231.

3. Gijsberts CM, Gohar A, Ellenbroek GH, Hoefer IE, de Kleijn DP, Asselbergs FW, Nathoe HM, Agostoni P, Rittersma SZ, Pasterkamp G, Appelman Y, den Ruijter HM. Severity of stable coronary artery disease and its biomarkers differ between men and women undergoing angiography. Atherosclerosis. 2015 Jul;241(1):234-40.

**VUNTR**

Since 1987, the Netherlands Twin Register is collecting (longitudinal) data in young and adult twins and their families [1,2]. The rich phenotypic longitudinal information that has been collected extends from lifestyle, exposures, personality and demographics information to mental and somatic health. In subgroups information on autonomic and central nervous system function, biomarkers and gene expression, epigenetics and genotyping is available. A 2015 estimate is that, since initiating the NTR, ~25% of all twins and multiples in the Netherlands participated in NTR research projects. Longitudinal information for over 200,000 participants (twins, multiples and family members) was collected over multiple NTR research projects. Data collection is ongoing. A pdf of nearly all published papers may be found at the NTR website.

As part of a Netherlands Twin Register (NTR) biobank project (BB1), 9,530 participants from 3,477 families were visited at home between January 2004 and July 2008 for collection of blood samples [3]. A second project (BB2) collected blood samples in 517 subjects from January 2011 to December 2011, including 210 MZ twin pairs and 64 twin-spouse pairs [4]. Visits were scheduled between 7:00 and 10:00 am and fertile women were bled on day 2-4 of the menstrual cycle, or in their pill-free week. Body composition was measured and information about physical health and lifestyle (e.g. smoking and drinking behavior, exercise, medication use) was obtained. For more detailed information about the methodology of the NTR Biobank study, see [3]. The NTR studies were approved by the Central Ethics Committee on Research involving human subjects of the VUMC, Amsterdam, an Institutional Review Board certified by the US Office of Human Research Protections (IRB number IRB-2991 under Federal wide Assurance-3703; IRB/institute codes, NTR 03-180). All subjects provided written informed consent. Subject were selected who were part of NTR biobank and who in general had rich phenotyping data available.

1. Van Beijsterveldt CE, Groen-Blokhuis M, Hottenga JJ, Franić S, Hudziak JJ, Lamb D, Huppertz C, de Zeeuw E, Nivard M, Schutte N, Swagerman S, Glasner T, van Fulpen M, Brouwer C, Stroet T, Nowotny D, Ehli EA, Davies GE, Scheet P, Orlebeke JF, Kan KJ, Smit D, Dolan CV, Middeldorp CM, de Geus EJ, Bartels M, Boomsma DI. The Young Netherlands Twin Register (YNTR): longitudinal twin and family studies in over 70,000 children. *Twin Res Hum Genet.* **2013** Feb; 16(1): 252-267. [doi: 10.1017/thg.2012.118](https://doi.org/10.1017/thg.2012.118). PMID: 23186620

2. Willemsen G, Vink JM, Abdellaoui A, den Braber A, van Beek JH, Draisma HH, van Dongen J, van 't Ent D, Geels LM, van Lien R, Ligthart L, Kattenberg M, Mbarek H, de Moor MH, Neijts M, Pool R, Stroo N, Kluft C, Suchiman HE, Slagboom PE, de Geus EJ, Boomsma DI.The Adult Netherlands Twin Register: twenty-five years of survey and biological data collection.*Twin Res Hum Genet.* **2013** Feb; 16(1): 271-281. [doi: 10.1017/thg.2012.140](https://doi.org/10.1017/thg.2012.140). PMID: 23298648. PMCID: PMC3739974.

3. Willemsen G, de Geus EJ, Bartels M, van Beijsterveldt CE, Brooks AI, Estourgie-van Burk GF, Fugman DA, Hoekstra C, Hottenga JJ, Kluft K, Meijer P, Montgomery GW, Rizzu P, Sondervan D, Smit AB, Spijker S, Suchiman HE, Tischfield JA, Lehner T, Slagboom PE, Boomsma DI. The Netherlands Twin Register biobank: a resource for genetic epidemiological studies. *Twin Res Hum Genet.* **2010** Jun; 13(3): 231-245. [doi: 10.1375/twin.13.3.231](https://doi.org/10.1375/twin.13.3.231). PMID: 20477721.

4. Sirota M, Willemsen G, Sundar P, Pitts SJ, Potluri S, Prifti E, Kennedy S, Ehrlich SD, Neuteboom J, Kluft C, Malone KE, Cox DR, de Geus EJ, Boomsma DI. Effect of genome and environment on metabolic and inflammatory profiles. *PLoS One.* **2015** Apr 8; 10(4): e012089. [doi: 10.1371/journal.pone.0120898](https://doi.org/10.1371/journal.pone.0120898). PMID: 25853885. PMCID: PMC4390246.

**website:** [www.tweelingenregister.org/](http://www.tweelingenregister.org/)
