## Supplemental Table S3 for "Predicting biological age based on the BBMRI-NL ^1^H-NMR metabolomics repository"

**Table S3. Description of metabolic variables in the predictor**

| **Metabolic variable** | **Description** | **Unit** |
| --- | --- | --- |
| **M-VLDL-L** | **Total lipids in medium VLDL** | **mmol/l** |
| **S-VLDL-L** | **Total lipids in small VLDL** | **mmol/l** |
| **XS-VLDL-L** | **Total lipids in very small VLDL** | **mmol/l** |
| **IDL-L** | **Total lipids in IDL** | **mmol/l** |
| **IDL-C** | **Total cholesterol in IDL** | **mmol/l** |
| **L-LDL-L** | **Total lipids in large LDL** | **mmol/l** |
| **M-LDL-L** | **Total lipids in medium LDL** | **mmol/l** |
| **M-HDL-L** | **Total lipids in medium HDL** | **mmol/l** |
| **S-HDL-L** | **Total lipids in small HDL** | **mmol/l** |
| **S-LDL-L** | **Total lipids in small LDL** | **mmol/l** |
| **VLDL-D** | **Mean diameter for VLDL particles** | **nm** |
| **LDL-D** | **Mean diameter for LDL particles** | **nm** |
| **HDL-D** | **Mean diameter for HDL particles** | **nm** |
| **Serum-C** | **Serum total cholesterol** | **mmol/l** |
| **VLDL-C** | **Total cholesterol in VLDL** | **mmol/l** |
| **LDL-C** | **Total cholesterol in LDL** | **mmol/l** |
| **HDL-C** | **Total cholesterol in HDL** | **mmol/l** |
| **HDL2-C** | **Total cholesterol in HDL2** | **mmol/l** |
| **HDL3-C** | **Total cholesterol in HDL3** | **mmol/l** |
| **Serum-TG** | **Serum total triglycerides** | **mmol/l** |
| **TotPG** | **Total phosphoglycerides** | **mmol/l** |
| **PC** | **Phosphatidylcholine and other cholines** | **mmol/l** |
| **SM** | **Sphingomyelins** | **mmol/l** |
| **TotCho** | **Total cholines** | **mmol/l** |
| **ApoA1** | **Apolipoprotein A-I** | **g/l** |
| **ApoB** | **Apolipoprotein B** | **g/l** |
| **TotFA** | **Total fatty acids** | **mmol/l** |
| **UnsatDeg** | **Estimated degree of unsaturation** |  |
| **DHA** | **22:6, docosahexaenoic acid** | **mmol/l** |
| **LA** | **18:2, linoleic acid** | **mmol/l** |
| **FAw3** | **Omega-3 fatty acids** | **mmol/l** |
| **FAw6** | **Omega-6 fatty acids** | **mmol/l** |
| **PUFA** | **Polyunsaturated fatty acids** | **mmol/l** |
| **MUFA** | **Monounsaturated fatty acids; 16:1, 18:1** | **mmol/l** |
| **SFA** | **Saturated fatty acids** | **mmol/l** |
| **FAw3/FA** | **Ratio of omega-3 fatty acids to total fatty acids** | **%** |
| **FAw6/FA** | **Ratio of omega-6 fatty acids to total fatty acids** | **%** |
| **PUFA/FA** | **Ratio of polyunsaturated fatty acids to total fatty acids** | **%** |
| **MUFA/FA** | **Ratio of monounsaturated fatty acids to total fatty acids** | **%** |
| **SFA/FA** | **Ratio of saturated fatty acids to total fatty acids** | **%** |
| **Glc** | **Glucose** | **mmol/l** |
| **Lac** | **Lactate** | **mmol/l** |
| **Cit** | **Citrate** | **mmol/l** |
| **Ala** | **Alanine** | **mmol/l** |
| **Gln** | **Glutamine** | **mmol/l** |
| **His** | **Histidine** | **mmol/l** |
| **Ile** | **Isoleucine** | **mmol/l** |
| **Leu** | **Leucine** | **mmol/l** |
| **Val** | **Valine** | **mmol/l** |
| **Phe** | **Phenylalanine** | **mmol/l** |
| **Tyr** | **Tyrosine** | **mmol/l** |
| **Ace** | **Acetate** | **mmol/l** |
| **AcAce** | **Acetoacetate** | **mmol/l** |
| **Crea** | **Creatinine** | **mmol/l** |
| **Alb** | **Albumin** | **signal area** |
| **GlycA** | **Glycoprotein acetyls, mainly a1-acid glycoprotein** | **mmol/l** |
| **bOHBut** | **3-hydroxybutyrate** | **mmol/l** |
| **XXL_VLDL_L** | **Total lipids in chylomicrons and extremely large VLDL** | **mmol/l** |
| **XL_VLDL_L** | **Total lipids in very large VLDL** | **mmol/l** |
| **L_VLDL_L** | **Total lipids in large VLDL** | **mmol/l** |
| **XL_HDL_L** | **Total lipids in very large HDL** | **mmol/l** |
| **L_HDL_L** | **Total lipids in large HDL** | **mmol/l** |

**In grey: 6 metabolomic variables measured at low success rates (<98%) or that frequently failed to reach the detection limit (<99%) were excluded.**
