## Supplemental Table S4 for "Predicting biological age based on the BBMRI-NL ^1^H-NMR metabolomics repository"

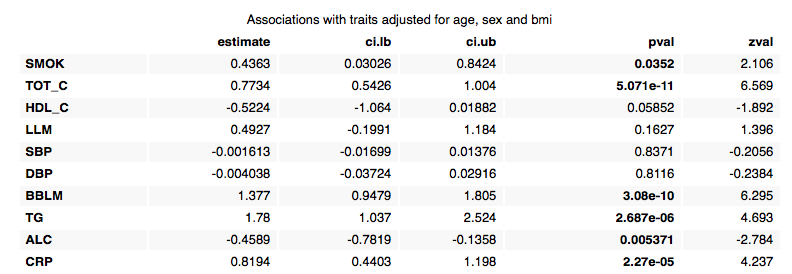

**Table S4**: *Associations with cardio-metabolic risk factors with ΔmetaboAge in BBMRI-NL adjusted for sex and BMI. SMOK: smoking [Y/N], TOT_C: Total cholesterol, HDL_C: HDL Cholesterol, LLM: Lipid Lowering Medicine, SBP: Systolic Blood Pressure, DBP: Diastolic Blood Pressure, BBLM: Blood Pressure Lowering Medication, TG: log(Triglycerides), ALC: Alcohol usage [Y/N] CRP: log(hsCRP). CI: confidence interval. P-values are in bold when nominal significant p≤0.05*.
