## Supplemental Table S5 for "Predicting biological age based on the BBMRI-NL ^1^H-NMR metabolomics repository"

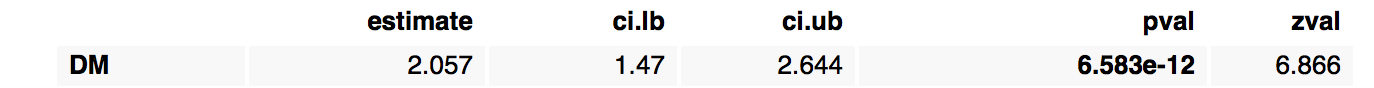

**TABLE S5:** *Associations of T2D status with ΔmetaboAge in BBMRI-NL adjusted for sex and BMI. CI: confidence interval. P-values are in bold when nominal significant p≤0.05.*
