## Supplemental Table S6 for "Predicting biological age based on the BBMRI-NL ^1^H-NMR metabolomics repository"

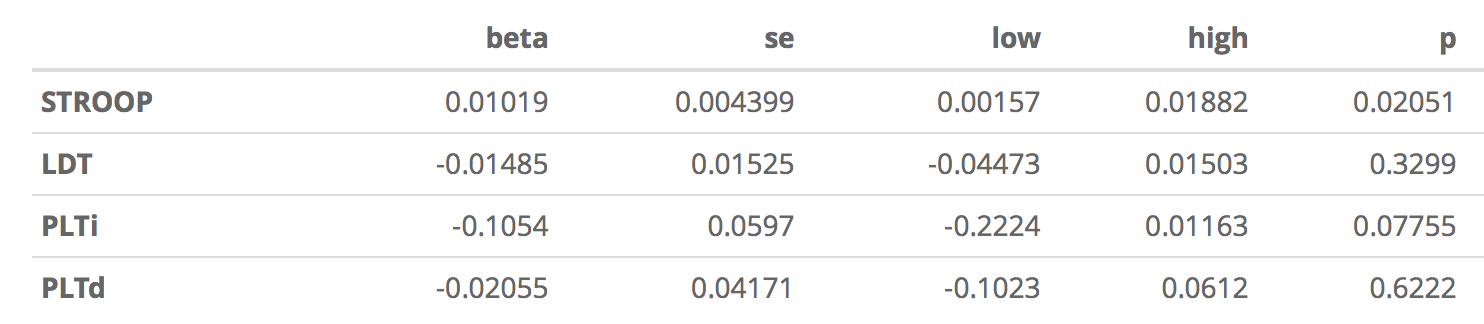

**Table S6:** *Associations of cognitive measures with ΔmetaboAge in PROSPER adjusted for sex, BMI, current smoking status [YES/NO], diabetes status [YES/NO] and hypertension status [YES/NO] and pravastatin treatment [YES/NO]. A small though significant effect is observed for the Stroop test that does not survive adjustments for multiple testing.*
