## Supplemental Table S7 for "Predicting biological age based on the BBMRI-NL ^1^H-NMR metabolomics repository"

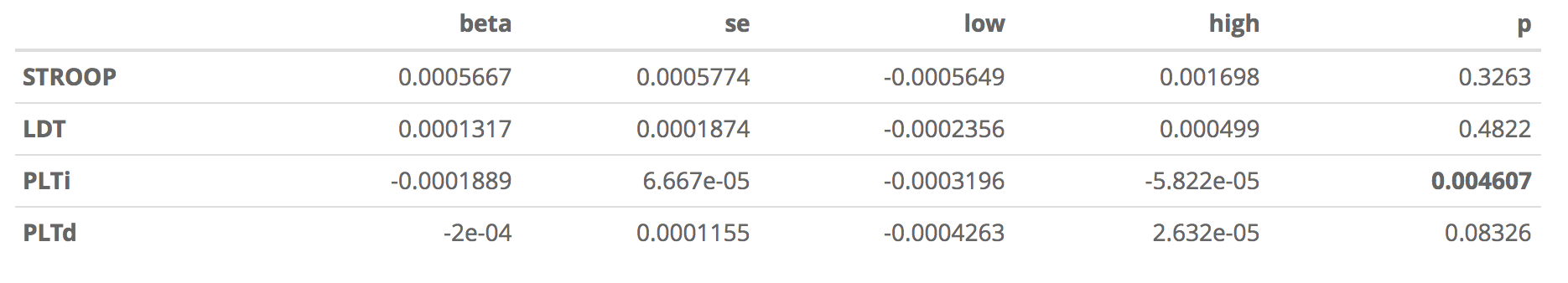

**Table S7:** *Associations of change in cognitive measures with ΔmetaboAge in PROSPER adjusted for sex, BMI, current smoking status [YES/NO], diabetes status [YES/NO] and hypertension status [YES/NO] and pravastatin treatment [YES/NO]. A significant though very marginal effect was observed for the change in PLTi.*
