## Supplemental Table S8 for "Predicting biological age based on the BBMRI-NL ^1^H-NMR metabolomics repository"

| **Phenotype** | **HR** | **95% CI** | ***p*-value** | **N_case_** | **N_total_** |
| --- | --- | --- | --- | --- | --- |
| Mortality | 1.16 | 1.06 – 1.26 | **0.001** | 744 | 761 |

**Table S8**: *Association of mortality with ΔmetaboAge in LLS_SIBS, adjusted for sex and IADL. HR reported as increased risk per 10-years of ΔmetaboAge.*
