## Supplemental Table S9 for "Predicting biological age based on the BBMRI-NL ^1^H-NMR metabolomics repository"

**Table S9**: Acknowledgements

**Alpha Omega**

Financial support for the Alpha Omega Cohort was obtained from the Dutch Heart Foundation (grant 200T401) and the National Institutes of Health (grant R01HL076200), USA. DNA isolation was funded by BBMRI-NL (grant CP2011-18). E. Waterham is acknowledged for data management and governance of the biobank of the Alpha Omega Cohort.

**Amsterdam Dementia Cohort**

Research within the VUmc Alzheimer center is part of the neurodegeneration research program of Amsterdam Neuroscience supported by Alzheimer Nederland and Stichting VUmc fonds.

**BIOMARCS**

**LUMINA**

LUMINA is supported by grants obtained from the Netherlands Organization for the Health Research and Development (ZonMw; no. 90700217) and VIDI (ZonMw; no. 91711319); the Netherlands Organisation for Scientific Research (NWO) VICI (no.918.56.602) and Spinoza prize (2009) grants; the Centre for Medical Systems Biology (CMSB) and Netherlands Consortium for Systems Biology (NCSB), both within the framework of the Netherlands Genomics Initiative (NGI)/Netherlands Organization for Scientific Research (NWO) the 7th Framework EU project EUROHEADPAIN (no. 602633).

**CHECK**

CHECK was funded by the Dutch Arthritis Association on the lead of a steering committee comprising 16 members with expertise in different fields of OA, chaired by Professor JWJ Bijlsma and coordinated by J Wesseling. Involved are: Erasmus Medical Center Rotterdam; Kennemer Gasthuis Haarlem; Leiden University Medical Center; Maastricht University Medical Center; Martini Hospital Groningen/Allied Health Care Center for Rheumatology and Rehabilitation Groningen; Medical Spectrum Twente Enschede/Ziekenhuisgroep Twente Almelo; Reade, formerly Jan van Breemen Institute/VU Medical Center Amsterdam; St. Maartenskliniek Nijmegen; University Medical Center Utrecht, and Wilhelmina Hospital Assen.

**CODAM**

Part of this work was supported by grants of the Netherlands Organisation for Scientific Research (940-35-034) and the Dutch Diabetes Research Foundation (98.901).

**The Maastricht Study (TMS)**

This study was supported by the European Regional Development Fund via OP-Zuid, the Province of Limburg, the Dutch Ministry of Economic Affairs (grant 31O.041), Stichting De Weijerhorst (Maastricht, the Netherlands), the Pearl String Initiative Diabetes (Amsterdam, the Netherlands), CARIM School for Cardiovascular Diseases (Maastricht, the Netherlands), Stichting Annadal (Maastricht, the Netherlands), Health Foundation Limburg (Maastricht, the Netherlands) and by unrestricted grants from Janssen-Cilag B.V. (Tilburg, the Netherlands), Novo Nordisk Farma B.V. (Alphen aan den Rijn, the Netherlands) and Sanofi-Aventis Netherlands B.V. (Gouda, the Netherlands).

**The Hoorn Diabetes Care System cohort study (DCS)**

This study was been made possible by collaboration with the Diabetes Care System West-Friesland. The authors thank participants of this study and research staff of the Diabetes Care System West-Friesland.

**Erasmus Rucphen Family (ERF) study**

The Erasmus Rucphen Family (ERF) study has received funding from the Centre for Medical Systems Biology (CMSB) and Netherlands Consortium for Systems Biology (NCSB), both within the framework of the Netherlands Genomics Initiative (NGI)/Netherlands Organization for Scientific Research (NWO). ERF study is also a part of EUROSPAN (European Special Populations Research Network) (FP6 STREP grant number 18947 (LSHG-CT-2006-018947)); European Network of Genomic and Genetic Epidemiology (ENGAGE) from the European Community's Seventh Framework Programme (FP7/2007-2013)/grant agreement HEALTH-F4-2007-201413; "Quality of Life and Management of the Living Resources" of 5th Framework Programme (no. QLG2-CT-2002-01254); FP7 project EUROHEADPAIN (nr 602633), the Internationale Stichting Alzheimer Onderzoek (ISAO); the Hersenstichting Nederland (HSN); and the JNPD under the project PERADES (grant number 733051021, Defining Genetic, Polygenic and Environmental Risk for Alzheimer’s Disease using multiple powerful cohorts, focused Epigenetics and Stem cell metabolomics). Metabolomics measurements of ERF has been funded by Biobanking and Biomolecular Resources Research Infrastructure (BBMRI)-NL (184.021.007). The ERF-follow up study is funded by CardioVasculair Onderzoek Nederland (CVON 2012-03). We are grateful to all study participants and their relatives, general practitioners and neurologists for their contributions and to P. Veraart for her help in genealogy, J. Vergeer for the supervision of the laboratory work, both S.J. van der Lee and A. van der Spek for collection of the follow-up data and P. Snijders for his help in data collection of both baseline and follow-up data.

**Rotterdam Study**

The Rotterdam Study is supported by the Erasmus MC University Medical Center and Erasmus University Rotterdam; The Netherlands Organisation for Scientific Research (NWO); The Netherlands Organisation for Health Research and Development (ZonMw); the Research Institute for Diseases in the Elderly (RIDE); The Netherlands Genomics Initiative (NGI); the Ministry of Education, Culture and Science; the Ministry of Health, Welfare and Sports; the European Commission (DG XII); and the Municipality of Rotterdam. The authors are grateful to the study participants, the staff from the Rotterdam Study and the participating general practitioners and pharmacists. Metabolomics measurements were funded by Biobanking and Biomolecular Resources Research Infrastructure (BBMRI)–NL (184.021.007) and the JNPD under the project PERADES (grant number 733051021, Defining Genetic, Polygenic and Environmental Risk for Alzheimer’s Disease using multiple powerful cohorts, focused Epigenetics and Stem cell metabolomics).

**FUNCTGENOMICS**

**GARP**

The Leiden University Medical Centre have and are supporting the RAAK and GARP study. This study was supported by the Dutch Arthritis Foundation and Pfizer Groton, Connecticut, USA. We are indebted to drs. N. Riyazi, J. Bijsterbosch, H.M. Kroon and I. Watt for collection of data.

**HELIUS**

The HELIUS study is conducted by the Academic Medical Center Amsterdam and the Public Health Service of Amsterdam. Both organisations provided core support for HELIUS. The HELIUS study is also funded by the Dutch Heart Foundation, the Netherlands Organization for Health Research and Development (ZonMw), the European Union (FP-7), and the European Fund for the Integration of non-EU immigrants (EIF).

**LIFELINES-DEEP**

**Leiden Longevity Study (LLS)**

The LLS has received funding from the European Union's Seventh Framework Programme (FP7/2007-2011) under grant agreement n° 259679. This study was supported by a grant from the Innovation-Oriented Research Program on Genomics (SenterNovem IGE05007), the Centre for Medical Systems Biology, and the Netherlands Consortium for Healthy Ageing (grants 05040202 and 050-060-810), all in the framework of the Netherlands Genomics Initiative, Netherlands Organization for Scientific Research (NWO), Unilever Colworth, and by BBMRI-NL, a Research Infrastructure financed by the Dutch government (NWO 184.021.007).

**NESDA**

The infrastructure for the NESDA study ([www.nesda.nl](http://www.nesda.nl/)) is funded through the Geestkracht program of the Netherlands Organisation for Health Research and Development (ZonMw, grant number 10-000-1002) and financial contributions by participating universities and mental health care organizations (VU University Medical Center, GGZ inGeest, Leiden University Medical Center, Leiden University, GGZ Rivierduinen, University Medical Center Groningen, University of Groningen, Lentis, GGZ Friesland, GGZ Drenthe, Rob Giel Onderzoekscentrum).

**PROspective Study of Pravastatin in the Elderly at Risk (PROSPER)**

The PROSPER study was supported by an investigator initiated grant obtained from Bristol-Myers Squibb. Prof. Dr. J. W. Jukema is an Established Clinical Investigator of the Netherlands Heart Foundation (grant 2001 D 032). PROSPER was supported by the European Federation of Pharmaceutical Industries Associations (EFPIA), Innovative Medicines Initiative Joint undertaking, European Medical Information Framework (EMIF) grant number 115372 and the European Commission under the Health Cooperation Work Programme of the 7th Framework Programme (Grant number 305507) “Heart ‘omics’ in AGEing” (HOMAGE).

**The LUMC arthroplasty studies**

This was a combination of TACTICS, TOMAAT and RAAK cohorts. TACTICS was funded by The Dutch Board of Health Care Insurances (College voor Zorgverzekeringen; OG99/023) and Sanquin Blood Bank. Involved were Prof. dr R.G.H.H. Nelissen, MD, Prof. dr A. Brand, MD, Leiden University Medical Centre; R.L. te Slaa MD, Reinier de Graaf Gasthuis, Delft; Dr R.G. Poll MD, Slotervaart ziekenhuis, Amsterdam; Dr K.M. Veenstra Franciscus ziekenhuis, Rotterdam and Prof. dr D. van Rhenen Sanquin Blood Bank, Rotterdam. Funding for the TOMAAT-study was received from ZonMW (06-601) and Sanquin Blood Supply (03-002), the Netherlands. Clinical Trial Number: ISRCTN96327523 (controlled-trials.com) and NTR 303 (Dutch Trial Register). The RAAK study was supported by the Leiden University Medical Centre. Furthermore, the molecular studies performed within to the RAAK study has received funding from the Dutch Arthritis Association (DAA_10_1-402), Biobanking and BioMolecular resources Research Infrastructure The Netherlands (BBMRI-NL) complementation project CP2013-84-CP2013-83 and Dutch Scientific Research council NWO /ZonMW VICI scheme (nr. 91816631/528).

**STEMI_GIPS-III**

**UCORBIO**

UCORBIO is conducted and supported by department of Cardiology, University Medical Center Utrecht, Netherlands. Folkert W. Asselbergs is supported by UCL Hospitals NIHR Biomedical Research Centre. Metabolic profiling was supported by Biobanking and BioMolecular resources Research Infrastructure, the Netherlands (BBMRI-NL). UCORBIO received funding from FP EU project CVgenes@target (HEALTH-F2-2013–601456). We would like to thank Ms. Jonne Hos and Ms. Merel Schurink for their logistical support and Daniel Kofink, PhD, for data management.

**VUNTR**

Funding was obtained from the Netherlands Organization for Scientific Research (NWO) and MagW/ZonMW grants 904-61-090, 985-10-002, 904-61-193,480-04-004, 400-05-717, Addiction-31160008, Middelgroot-911-09-032, Spinozapremie 56-464-14192, Biobanking and Biomolecular Resources Research Infrastructure (BBMRI –NL, 184.021.007).; the European Community's Seventh Framework Program (FP7/2007-2013), ENGAGE (HEALTH-F4-2007-201413); the European Science Council (ERC Advanced, 230374), Rutgers University Cell and DNA Repository (NIMH U24 MH068457-06), the Avera Institute, Sioux Falls, South Dakota (USA) and the National Institutes of Health (NIH, R01D0042157-01A, MH081802, Grand Opportunity grants 1RC2 MH089951). We gratefully acknowledge grant NWO 480-15-001/674: Netherlands Twin Registry Repository: researching the interplay between genome and environment.
