## Supplemental Material S1 for "Predicting biological age based on the BBMRI-NL ^1^H-NMR metabolomics repository"

### DocS1

*ebvdakker*

*2018-10-01*

```
## Warning: package 'knitr' was built under R version 3.4.3
```

```
## || Loading data:
## ../preprocessing/BBMRI__2018-03-15_23-00-34.308.RData
## | Excluding 5737 entries in 'PROSPER', 'LLS_SIBS', 'VUNTR', 'CODAM' ... CODAM [N=145]; LLS_SIBS [N=9
## | Removing 100 entries with missing age, sex or age<18  ... ALPHAOMEGA [N=1]; BIOMARCS [N=3]; CHARM
## | Removing discontinued metabolites 'dag', 'dag_tg', 'falen', 'cla', 'cla_fa' ... Done!
## | Removing metabolites not reported in serum 'pyr' ... Done!
## | Removing dependent metabolites 'apob_apoa1', 'dha_fa', 'estc', 'fast', 'freec', 'hdl_tg', 'idl_c_p
## | Removing metabolites showing many missing or zero values 'bohbut', 'xl_vldl_l', 'xxl_vldl_l', 'l_v
## | Removing 200 entries with missing values [Nmax>=1] ... ALPHAOMEGA [N=5]; BIOMARCS [N=62]; CHECK [N
## | Removing 64 entries with zero values [Nmax>=1] ... BIOMARCS [N=7]; CHECK [N=1]; CSF [N=1]; DCS [N=
## | Removing 442 entries with a 5SD outlier ... ALPHAOMEGA [N=92]; BIOMARCS [N=29]; CHARM [N=2]; CHECK
## | Scaling ... Done!
## | Imputing  467 [ 0.044 %] missing values ... Done!
```

### 5-Crossfold validation

#### Split train and test sets

```
##       train test
## Fold1 15208 3802
## Fold2 15208 3802
## Fold3 15208 3802
## Fold4 15209 3801
## Fold5 15207 3803
```

### Fit linear model on all train:

### Plot linear model fits

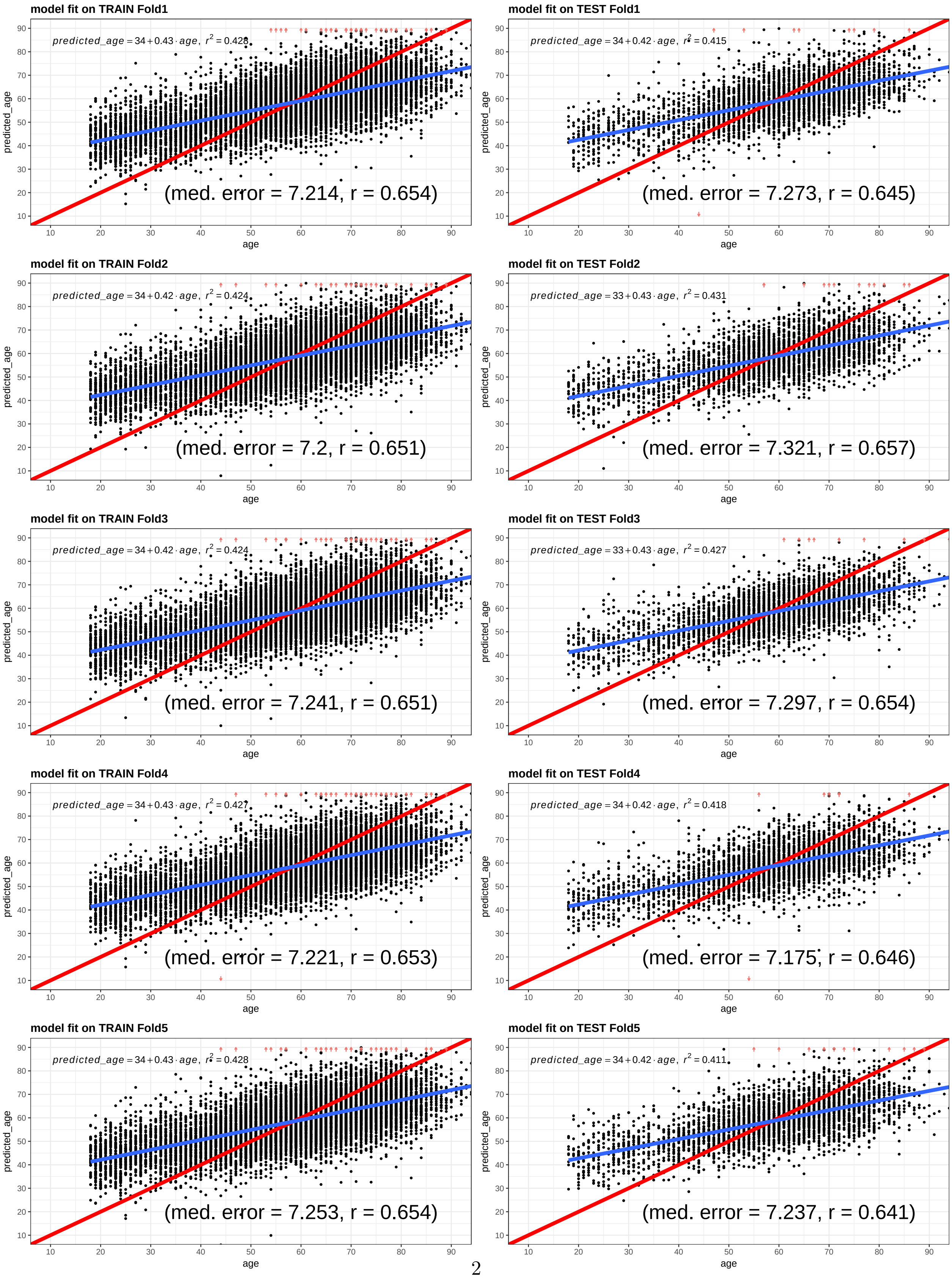

### Leave-One-Biobank-Out-Validation

#### Split train and test sets

```
##                train test
## ALPHAOMEGA     18231  779
## BIOMARCS       18270  740
## CHARM          18802  208
## CHECK          18045  965
## CSF            18756  254
## DCS            18054  956
## ERF            17624 1386
## FUNCTGENOMICS  18550  460
## GARP           18618  392
## HELIUS         18569  441
## LIFELINES      17552 1458
## LLS_PARTOFFS   16756 2254
## MRS            18921   89
## NESDA          17461 1549
## RAAK           18899  111
## RS             16083 2927
## STEMI_GIPS-III 18683  327
## TACTICS        18914   96
## TMS            18188  822
## TOMAAT         18785  225
## UCORBIO        17891 1119
## VUMC_ADC       17558 1452
```

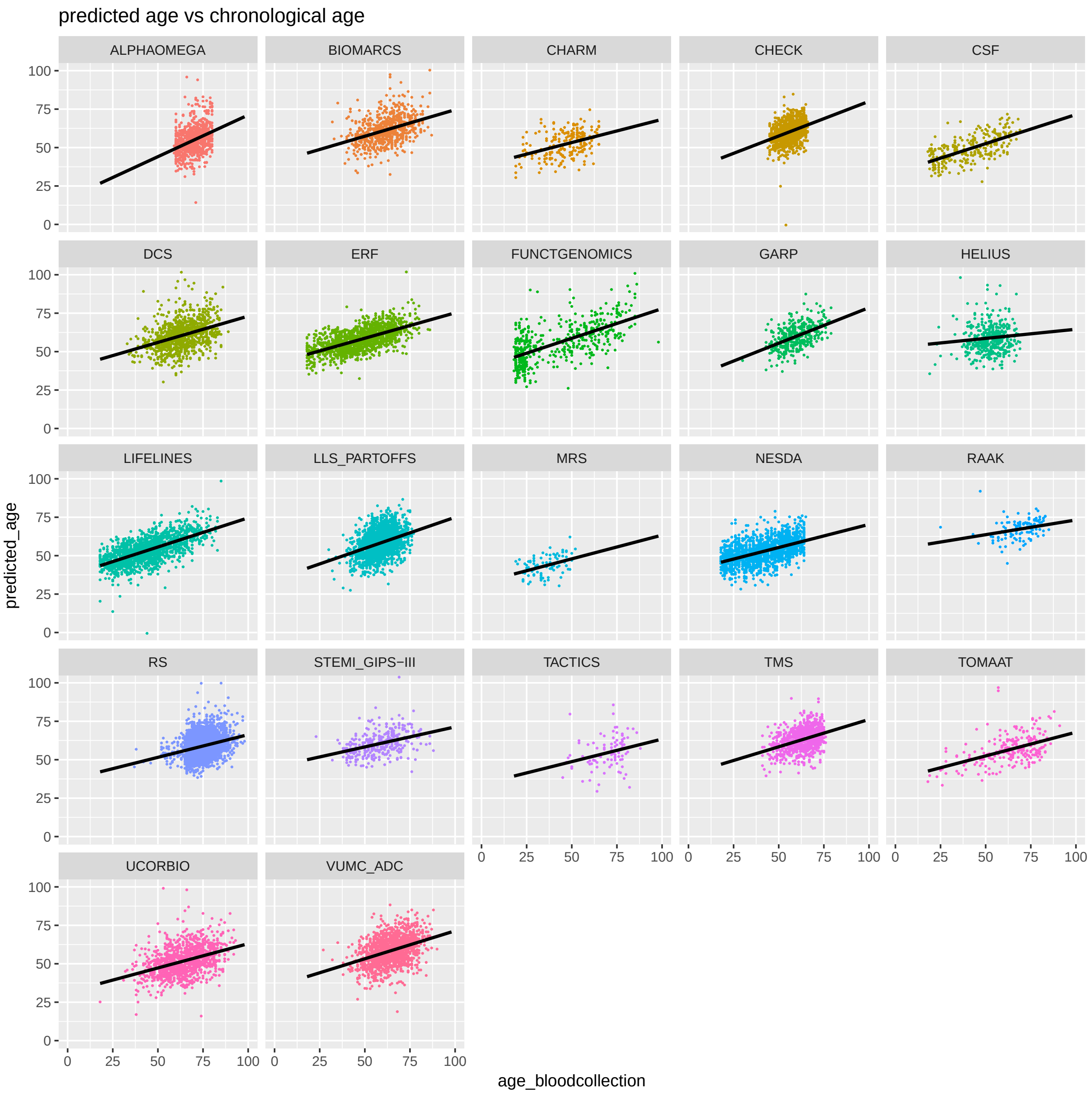
predicted age vs chronological age
ALPHAOMEGA
BIOMARCS
CHARM
CHECK
CSF
DCS
ERF
FUNCTGENOMICS
GARP
HELIUS
LIFELINES
LLS_PARTOFFS
MRS
NESDA
RAAK
RS
STEMI_GIPS−III
TACTICS
TMS
TOMAAT
UCORBIO
VUMC_ADC
predicted_age
age_bloodcollection
0
25
50
75
100
