## Supplemental Material S2 for "Predicting biological age based on the BBMRI-NL ^1^H-NMR metabolomics repository"

### DocS2 / QC REF

*ebvdakker*

*2018-10-01*

**Transform & Impute** **6**

**sessionInfo** **11**

```
## Warning: package 'knitr' was built under R version 3.4.3
```

### Load

```
## || Loading data:
## ../preprocessing/BBMRI__2018-03-15_23-00-34.308.RData
## | Removing 100 entries with missing age, sex or age<18  ... ALPHAOMEGA [N=1]; BIOMARCS [N=3]; CHARM
## | Removing discontinued metabolites 'dag', 'dag_tg', 'falen', 'cla', 'cla_fa' ... Done!
## | Removing metabolites not reported in serum 'pyr' ... Done!
## | Removing dependent metabolites 'apob_apoa1', 'dha_fa', 'estc', 'fast', 'freec', 'hdl_tg', 'idl_c_p
```

```
##         mat  phen
## [1,]     62 25453
## [2,] 25453    73
```

### QC per BIOBANK

#### Summary

```
##                   N Males          Age
## ALPHAOMEGA      876  77.1  69.7 [59-80]
## BIOMARCS        838  78.3  62.1 [32-87]
## CHARM           210  25.7  46.1 [19-65]
## CHECK           975  20.9  56.0 [44-66]
## CODAM           145  65.5  60.8 [44-72]
## CSF             259  39.8  40.9 [18-69]
```

```
## TMS               855  68.1  62.8 [41-76]
## DCS               995  57.1  62.9 [33-89]
## RS               2972  42.1  74.3 [37-98]
## ERF              1398  44.6  47.7 [18-86]
## FUNCTGENOMICS     497  64.8  42.2 [18-98]
## GARP              400  22.2  59.2 [30-79]
## HELIUS            472  41.3  52.1 [19-69]
## LIFELINES        1486  42.5  44.8 [18-85]
## LLS_PARTOFFS     2313  44.4  58.7 [30-80]
## LLS_SIBS          998  38.2 92.1  [53-103]
## MRS                90  50.0  35.2 [19-52]
## NESDA            1587  34.5  41.8 [18-65]
## PROSPER           983  51.4  75.6 [70-83]
## RAAK              121  28.1  69.1 [25-85]
## STEMI_GIPS-III    346  77.5  58.0 [23-88]
## TACTICS           115  18.3  69.7 [45-88]
## TOMAAT            233  37.8  63.5 [18-91]
## UCORBIO          1198  72.8  64.4 [18-93]
## VUMC_ADC         1480  55.6  64.0 [27-90]
##  [ reached getOption("max.print") -- omitted 1 row ]
```

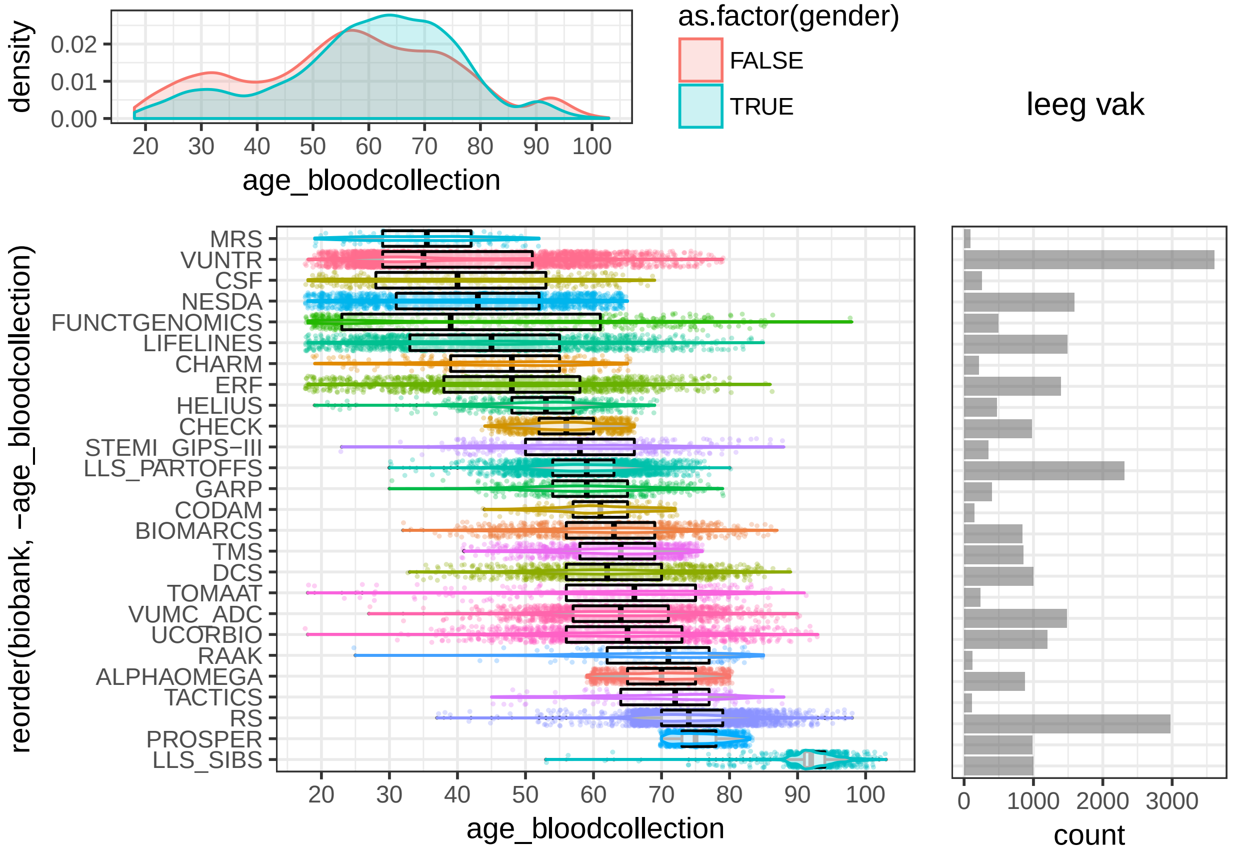

```
## TableGrob (4 x 4) "arrange": 4 grobs
##   z     cells    name           grob
## 1 1 (2-4,1-3) arrange gtable[layout]
## 2 2 (2-4,4-4) arrange gtable[layout]
## 3 3 (1-1,1-3) arrange gtable[layout]
## 4 4 (1-1,4-4) arrange text[GRID.text.1]
```

#### Holdout biobanks

Due to the age-range of **LLS_SIBS** and the additional samples available for **PROSPER** outside BBMRI, it was decided to use these as **HOLDOUT**.

```
## | Excluding 1981 entries in 'PROSPER', 'LLS_SIBS' ... LLS_SIBS [N=998]; PROSPER [N=983] ... Done!
```

#### Missingness per biobank

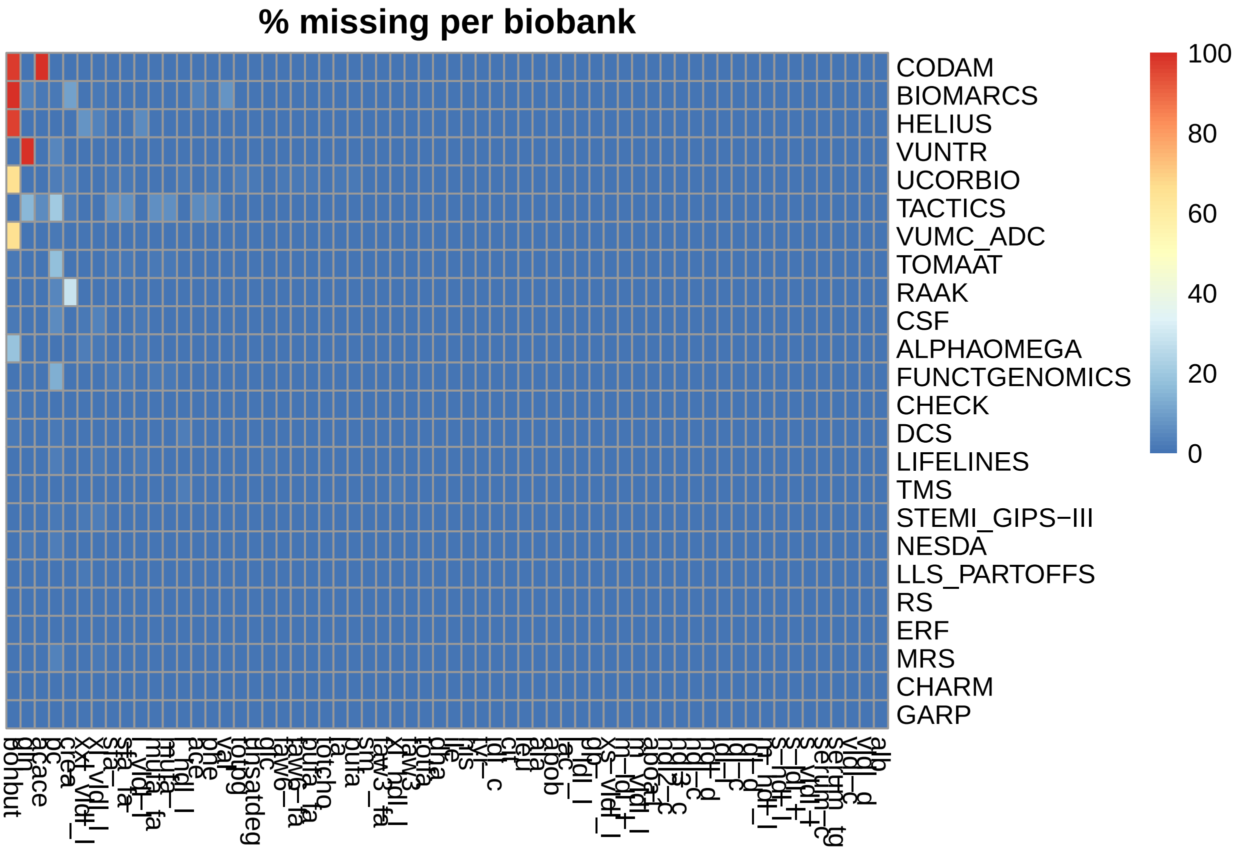

VUNTR and CODAM are uniquely fully missing a metabolite, respectively 'gln' and 'acace'. It was decided to remove these studies from the reference. The metabolite bohbut proved to be a problematic in multiple studies. Hence, it was decided to remove 'bohbut' from the reference.

```
## | Excluding 3756 entries in 'VUNTR', 'CODAM' ... CODAM [N=145]; VUNTR [N=3611] ... Done!
```

```
## | Removing 'bohbut' ... Done!
```

##### Zeros per biobank

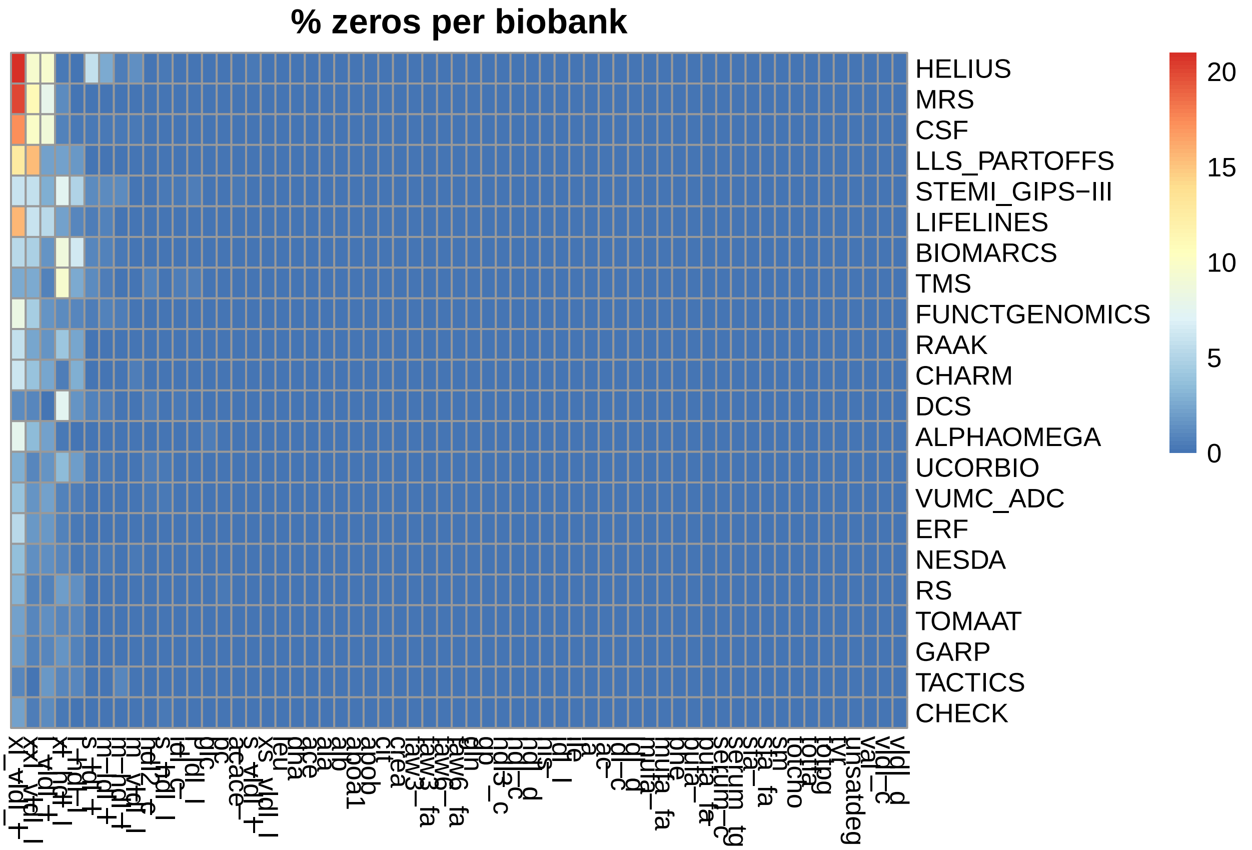

Metabolites frequently below the detection threshold in multiple biobanks were removed from the reference and included: xl__vldl__l, xxl__vldl__l, l__vldl__l, xl__hdl__l, l__hdl__l.

```
## | Removing 'xl_vldl_l', 'xxl_vldl_l', 'l_vldl_l', 'xl_hdl_l', 'l_hdl_l' ... Done!
```

#### QC per SAMPLE

##### Misingness per sample

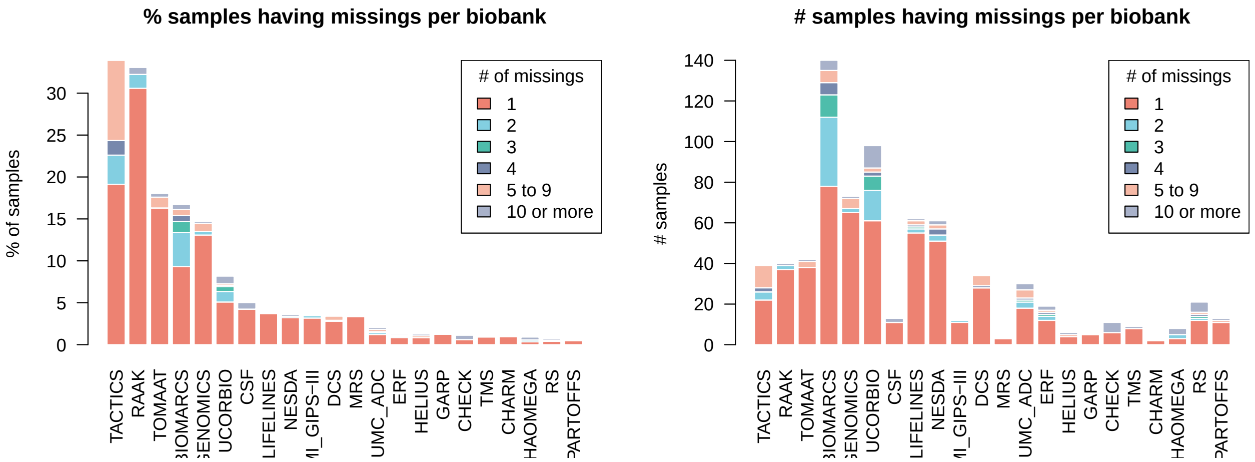

It was decided to remove samples having more than 1 missing value from the reference.

```
## | Removing 200 entries with missing values [Nmax>=1] ... ALPHAOMEGA [N=5]; BIOMARCS [N=62]; CHECK [N=
```

#### Zeros per sample

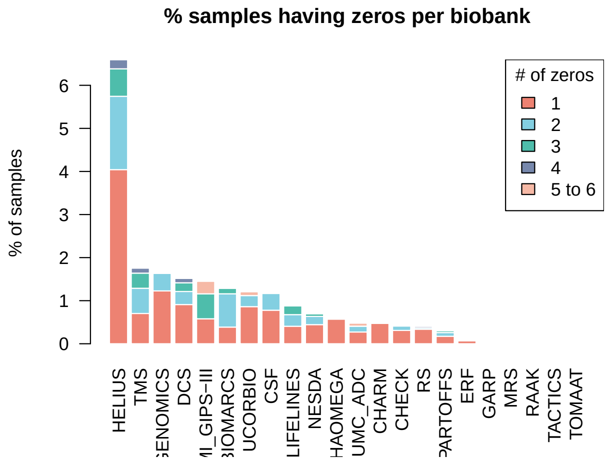

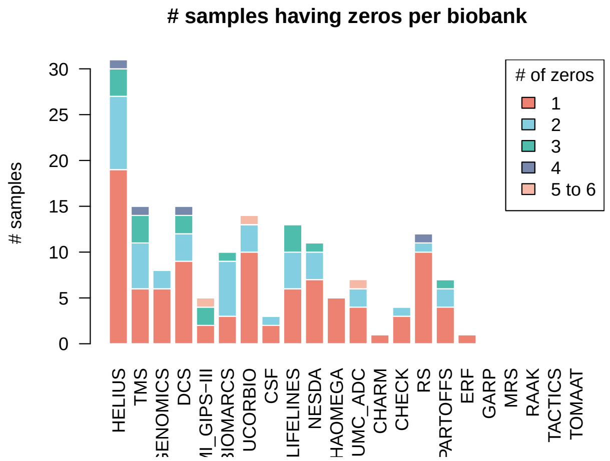

It was decided to remove samples having more than 1 zero value from the reference.

```
## | Removing 64 entries with zero values [Nmax>=1] ... BIOMARCS [N=7]; CHECK [N=1]; CSF [N=1]; DCS [N=
```

#### Cross-consortium 5SD Outliers

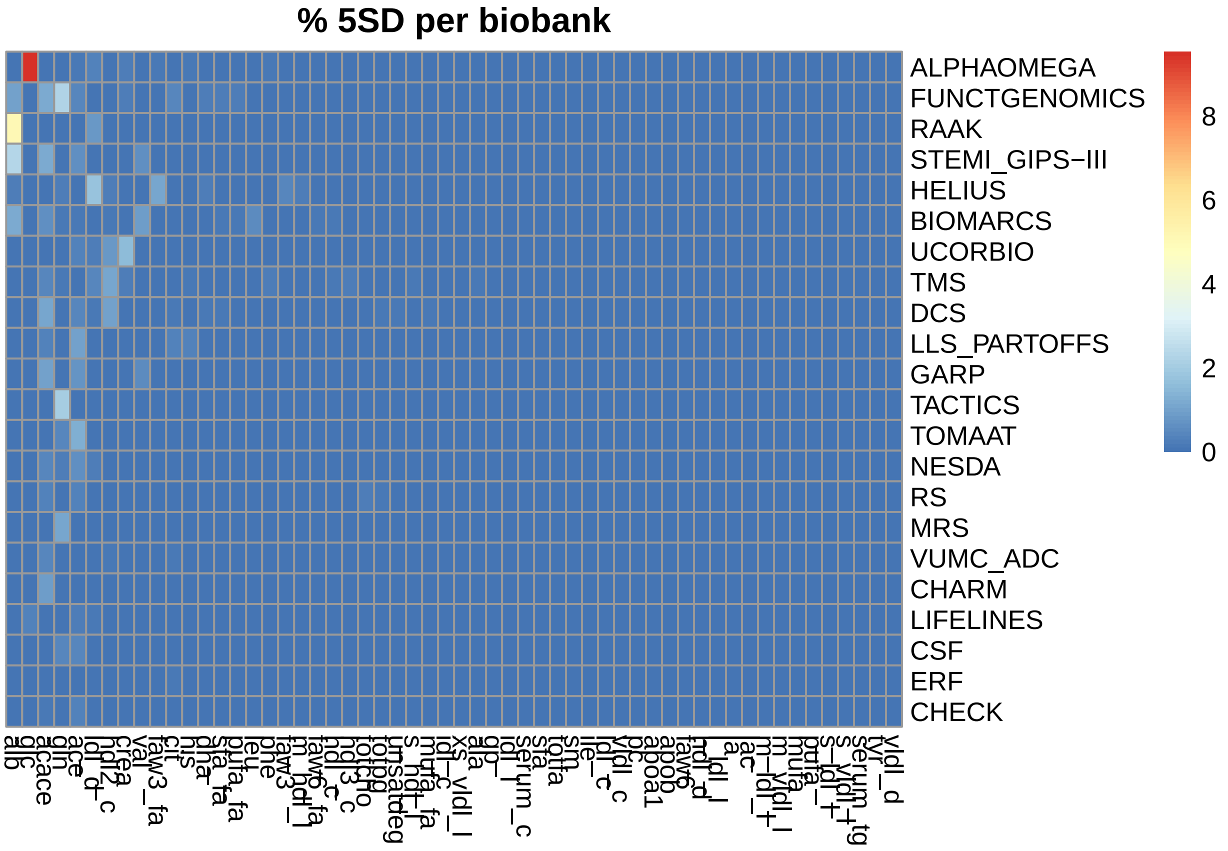

It was decided to remove all samples having a 5SD outlier value.

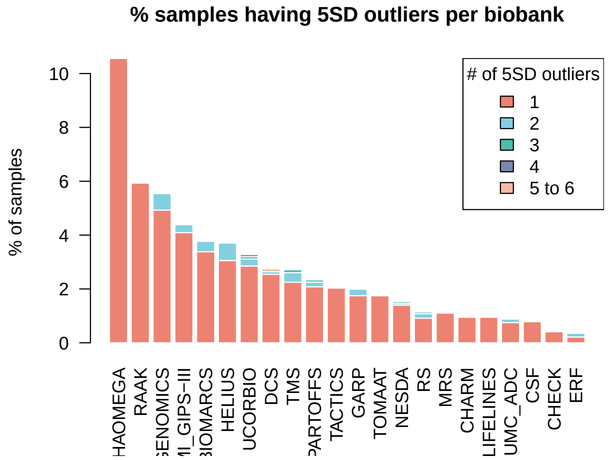

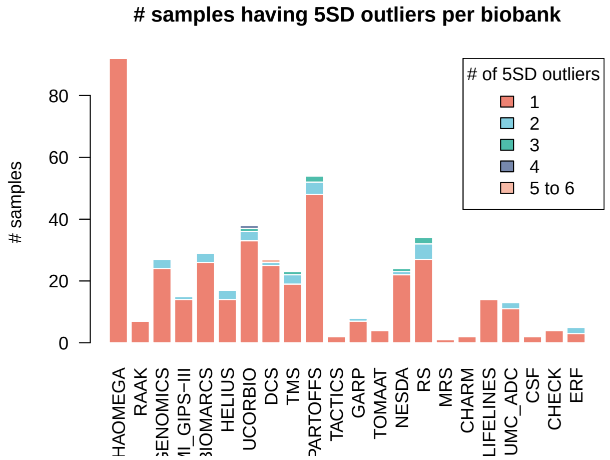

```
## | Removing 442 entries with a 5SD outlier ... ALPHAOMEGA [N=92]; BIOMARCS [N=29]; CHARM [N=2]; CHECK
```

### Transform & Impute

Since metabolite distributions generaly look reasonably gaussian shaped, only scaling is performed. Imputation of the remaining 467 datapoints is performed with 'nipals' imputation.

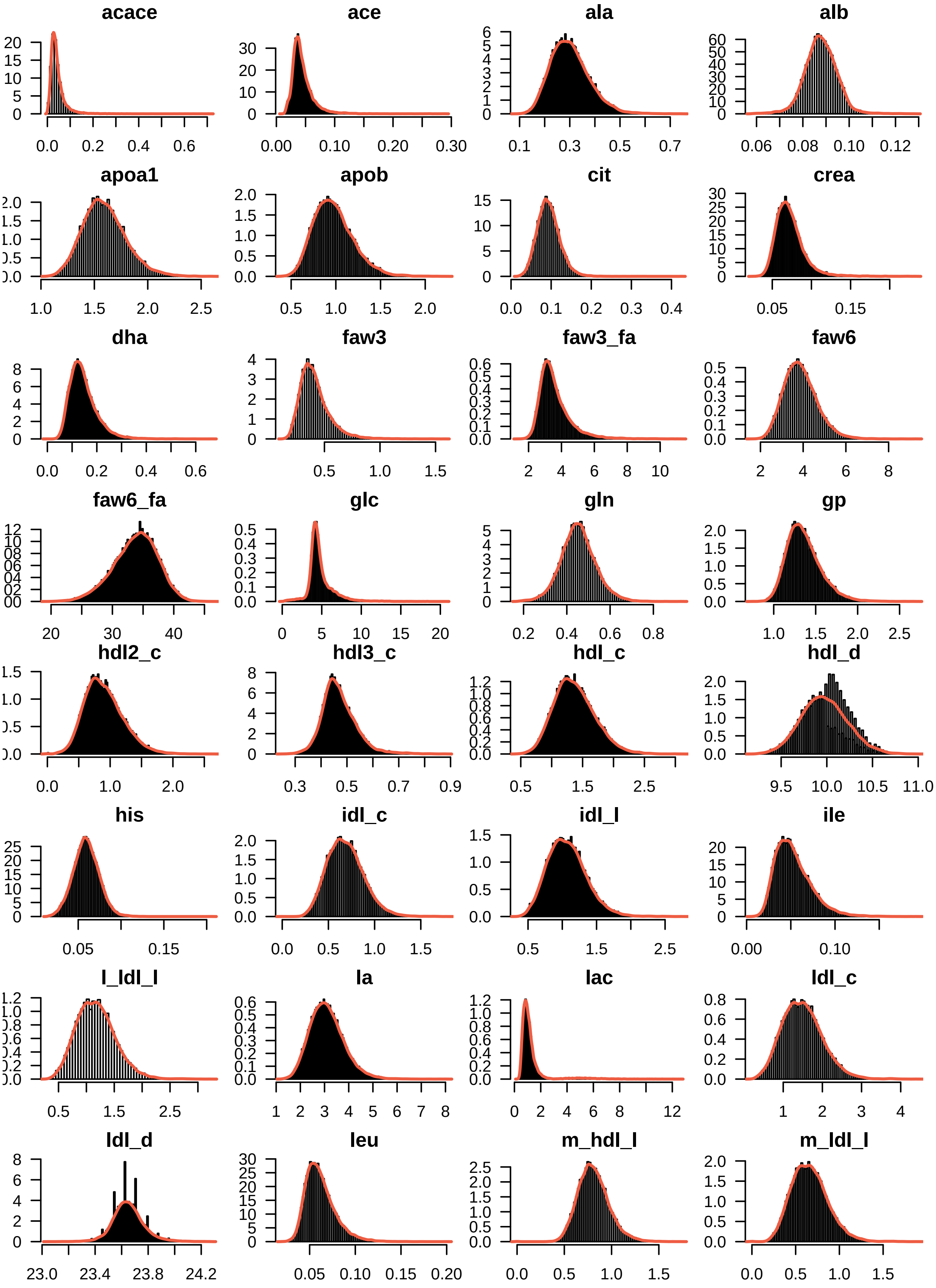
acace
0.0 0.2 0.4 0.6
ace
0.00 0.10 0.20 0.30
ala
0.1 0.3 0.5 0.7
alb
0.06 0.08 0.10 0.12
apoa1
1.0 1.5 2.0 2.5
apob
0.5 1.0 1.5 2.0
cit
0.0 0.1 0.2 0.3 0.4
crea
0.05 0.15
dha
0.0 0.2 0.4 0.6
faw3
0.5 1.0 1.5
faw3_fa
2 4 6 8 10
faw6
2 4 6 8
faw6_fa
20 30 40
glc
0 5 10 15 20
gln
0.2 0.4 0.6 0.8
gp
1.0 1.5 2.0 2.5
hdl2_c
0.0 1.0 2.0
hdl3_c
0.3 0.5 0.7 0.9
hdl_c
0.5 1.5 2.5
hdl_d
9.5 10.0 10.5 11.0
his
0.05 0.15
idl_c
0.0 0.5 1.0 1.5
idl_l
0.5 1.5 2.5
ile
0.00 0.10
l_ldl_l
0.5 1.5 2.5
la
1 2 3 4 5 6 7 8
lac
0 2 4 6 8 12
ldl_c
1 2 3 4
ldl_d
23.0 23.4 23.8 24.2
leu
0.05 0.10 0.15 0.20
m_hdl_l
0.0 0.5 1.0 1.5
m_ldl_l
0.0 0.5 1.0 1.5

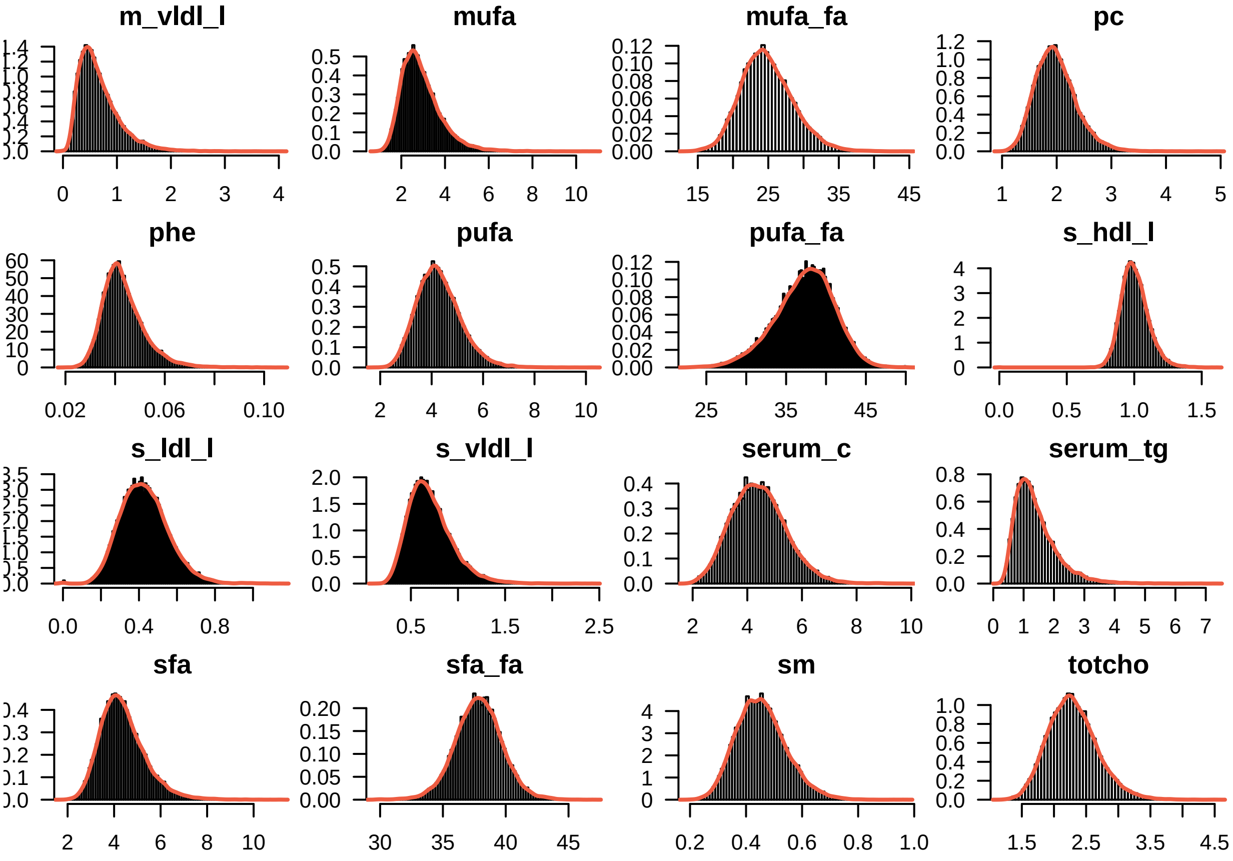

```
## | Scaling ... Done!
```

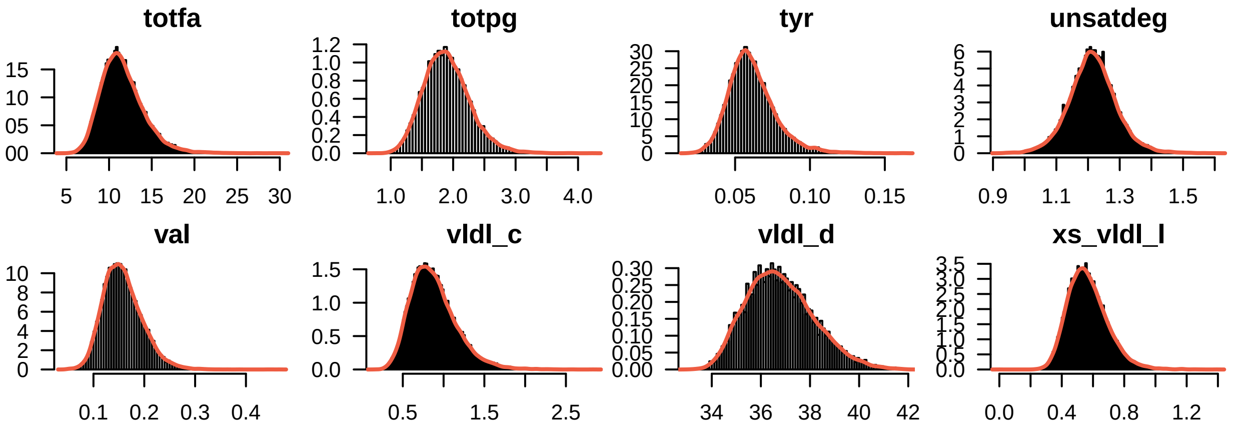

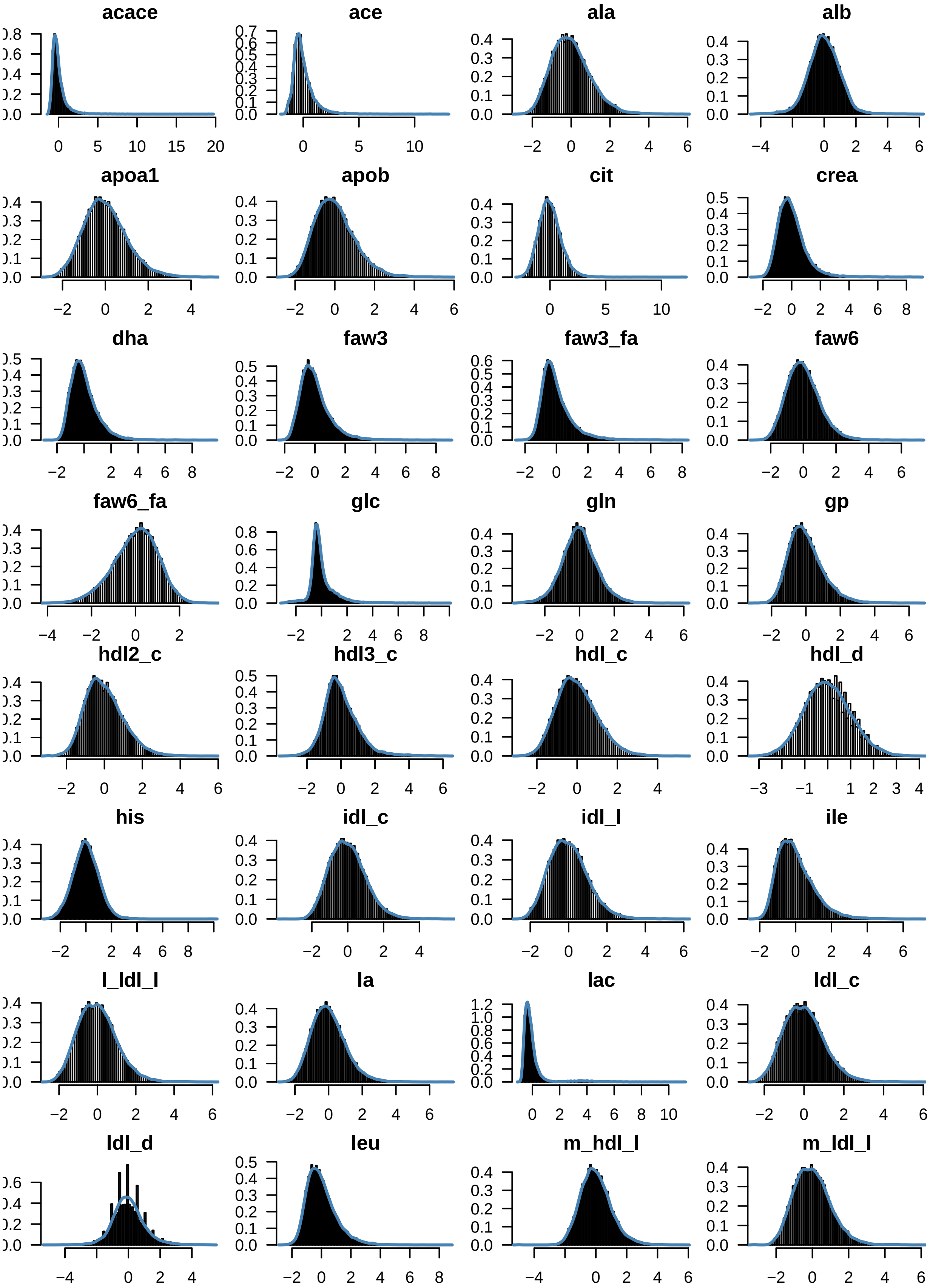
acace
ace
ala
alb
apoa1
apob
cit
crea
dha
faw3
faw3_fa
faw6
faw6_fa
glc
gln
gp
hdl2_c
hdl3_c
hdl_c
hdl_d
his
idl_c
idl_l
ile
l_ldl_l
la
lac
ldl_c
ldl_d
leu
m_hdl_l
m_ldl_l

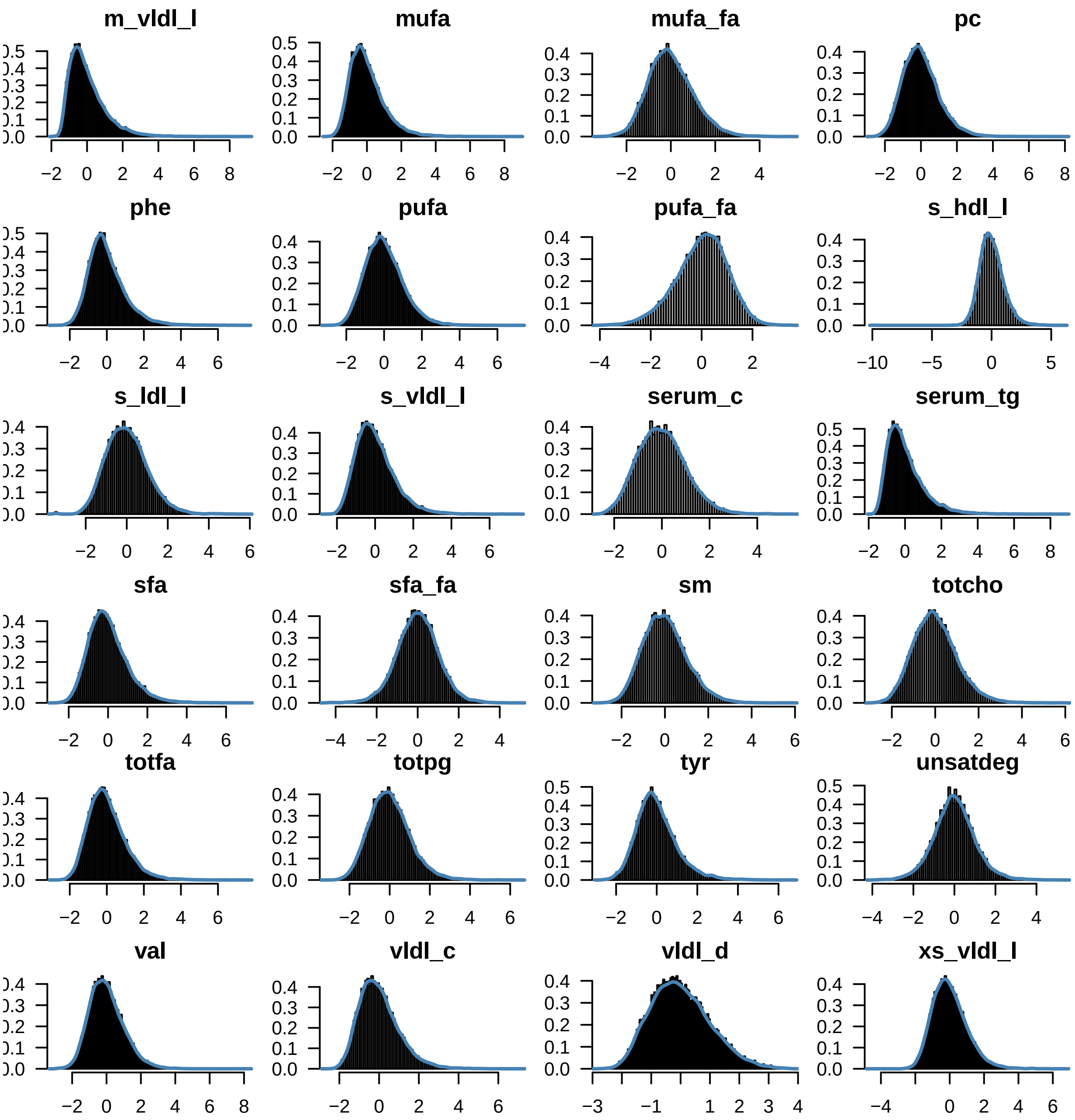

```
## | Imputing  467 [ 0.044 %] missing values ...
```

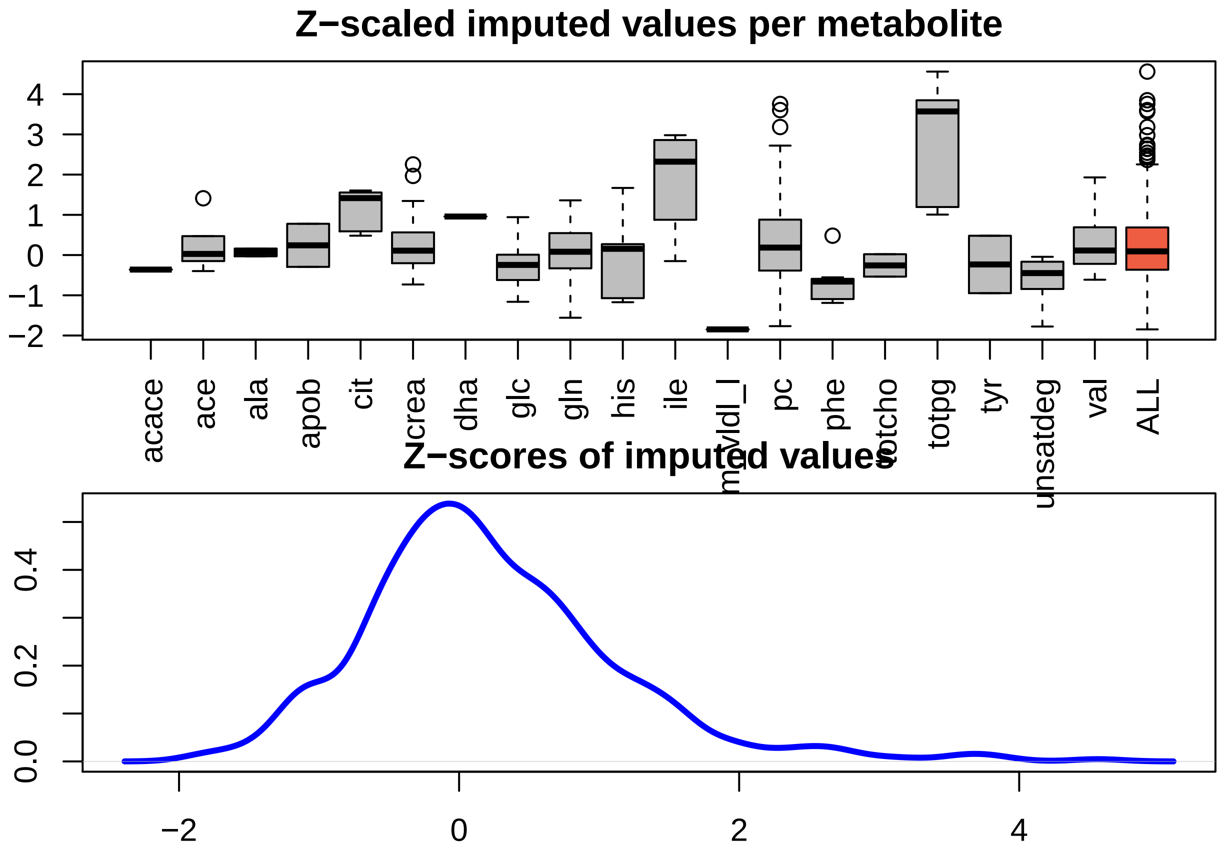

```
## Done!
```

#### sessionInfo

```
## R version 3.4.0 (2017-04-21)
## Platform: x86_64-apple-darwin15.6.0 (64-bit)
## Running under: OS X El Capitan 10.11.6
##
## Matrix products: default
## BLAS: /Library/Frameworks/R.framework/Versions/3.4/Resources/lib/libRblas.0.dylib
## LAPACK: /Library/Frameworks/R.framework/Versions/3.4/Resources/lib/libRlapack.dylib
##
## locale:
## [1] en_US.UTF-8/en_US.UTF-8/en_US.UTF-8/C/en_US.UTF-8/en_US.UTF-8
##
## attached base packages:
## [1] parallel  stats     graphics  grDevices utils     datasets  methods
## [8] base
##
## other attached packages:
## [1] ggsci_2.7           pcaMethods_1.68.0   Biobase_2.36.2
## [4] BiocGenerics_0.22.0 pheatmap_1.0.8      ggplot2_2.2.1
## [7] knitr_1.20
##
## loaded via a namespace (and not attached):
##  [1] Rcpp_0.12.16       magrittr_1.5       munsell_0.4.3
```

```
##  [4] colorspace_1.3-2   rlang_0.1.1        stringr_1.3.1
##  [7] plyr_1.8.4         tools_3.4.0        grid_3.4.0
## [10] gtable_0.2.0       htmltools_0.3.6    yaml_2.1.19
## [13] lazyeval_0.2.0     rprojroot_1.3-2    digest_0.6.12
## [16] tibble_1.3.4       gridExtra_2.2.1    RColorBrewer_1.1-2
## [19] evaluate_0.10.1    rmarkdown_1.10.2   labeling_0.3
## [22] stringi_1.2.2      compiler_3.4.0     scales_0.5.0
## [25] backports_1.1.2
```
